# Single-cell transcriptomics reveals the impact of sex and age in the healthy human liver

**DOI:** 10.1101/2025.01.27.635138

**Authors:** Raza Ur Rahman, Eliana T. Epstein, Shane Murphy, Liat Amir-Zilberstein, Cristin McCabe, Toni Delorey, Hope Koene, Lilly Fernandes, Kenneth K. Tanabe, Motaz Qadan, Christina Ferrone, David L. Berger, Angela Shih, Jacques Deguine, Alan C. Mullen

## Abstract

**Background & Aims:** The liver is a vital organ composed of parenchymal, nonparenchymal, and immune cell populations. Single-cell sequencing approaches now provide the opportunity to understand how sex and age influence gene expression and cellular function across cell types within the liver.

**Methods:** We analyzed the cellular composition and interactions for the human liver through single-nucleus RNA sequencing (snRNA-seq), incorporating insights from 37 healthy liver samples. The dataset contains cells from female and male donors spanning more than seven decades of life, and analysis was performed to evaluate the impact of sex and age on differential gene expression, pathway enrichment, and predicted ligand-receptor and protein-protein interactions.

**Results:** Excluding the X and Y chromosomes, we identified 374 genes uniquely enriched in cells of the female liver and 520 genes enriched in cells of the male liver. Differential expression analysis defined unique circuitries enriched within each cell type between females and males and their impact on cell-cell communication and response to external signals, including enrichment of cholesterol/lipid metabolism, transforming growth factor beta (TGF-beta) signaling, and fibronectin (FN1) production in female cells and bone morphogenic protein (BMP) signaling in male cells. With increased age, we observe a greater diversity in gene expression, including enrichment of genes that regulate neuregulin (NGR) signaling at older ages, while genes regulating insulin growth factor (IGF) signaling are enriched at younger ages. Senescence signatures were also identified for each cell type within the liver.

**Conclusions:** These results define the activities of healthy cell types within the liver across sex and age and provide a foundation for studies to examine how ancestry, geography, and disease states influence liver function within these contexts.

**Impact and Implications:** Our study analyzes 37 human liver samples at the single-cell level to understand how sex and age influence gene expression, cell interactions, and response to signals across liver cell types and sub-types. These findings are of particular significance for researchers who need to understand how sex and age may influence the response of individual cell types to injury or treatment of injury. This dataset will also provide a healthy reference for future studies to understand how ancestry, geography, and disease states shape liver biology across age and sex.

## INTRODUCTION

The liver plays a pivotal role in numerous physiological processes, including glucose and lipid metabolism, coagulation, iron homeostasis, bile secretion, and immune interactions. Dissecting the complex cellular composition of the liver and the influence of factors such as sex and age is essential to translate study findings in both health and disease to more specific patient populations. Advancements in single-cell RNA sequencing (scRNA-seq) technologies have provided unprecedented insights into the cellular landscape of the liver [1–4]. The introduction of single-nucleus RNA sequencing (snRNA-seq) has also allowed for broader representation of non-immune cell types in the liver compared to scRNA-seq [5], as these cell types may be more sensitive to enzymatic digestion and mechanical separation. While extensive research has been conducted to examine the liver at the single cell resolution [1–4], snRNA-seq data for human liver [5] remain limited, and larger datasets are necessary to understand the full extent to which factors such as sex and age affect the function of the diverse cell types and subtypes within the liver.

Sex-based variations in enzyme activity, gene expression, and steroid hormone sensitivity can influence the ability of the liver to metabolize drugs and hormones [6–8]. Chronic liver diseases such as fibrosis and cirrhosis are more prevalent and severe in men than in women [9–11], and for this reason male mice are used more frequently in models of *in vivo* liver injury [12–14]. While single cell analyses have defined the transcriptional activity of individual cell types, cell-cell interactions, and how these features change in disease [2,15–18], our understanding of sex-specific pathways at single-cell resolution remains limited.

Aging is also an established risk factor for chronic diseases, including those that involve the liver, and the global population is aging rapidly. In 2020, the number of people 60 years of age and older surpassed those under age 5, and the percentage of individuals 60 years of age and older is projected to nearly double from 2015 to 2050, increasing from 12% to 22% of the global population (https://www.who.int/news-room/fact-sheets/detail/ageing-and-health, October 2024).

Within the liver, aging leads to changes in gene expression, cell function, and morphology [19–24] and is linked to reduced regenerative capacity, increased susceptibility to fibrosis, and greater vulnerability to conditions including MASLD, metabolic-dysfunction associated steatohepatitis (MASH), and cirrhosis. Elderly patients with MASLD have a higher prevalence of inflammation/MASH and advanced fibrosis compared to younger counterparts [25]. One mechanism that may promote the increased prevalence of MASLD with age is cellular senescence, and clearance of senescent hepatocytes can reduce hepatic steatosis/MASLD in aging and obesity models [26]. Additional cell types, including endothelial cells and macrophages also show induction of senescence signatures with age [27]. A more complete understanding of the changes that occur in each cell type and subtype with age will provide insights into the broader implications of liver aging, response to injury, and potential interventions to mitigate its effects.

Despite significant progress in understanding the impact of sex and age on liver function and health, our knowledge at the single cell level remains limited. Here, we present a comprehensive map of the cellular composition of the human liver through snRNA-seq, analyzing data from 37 healthy liver samples (>195,000 nuclei) across seven decades of life from both female and male donors. This study highlights how sex and age influence gene expression, enhances our understanding of cellular heterogeneity within the liver, and provides a foundation for future research on sex- and age-related disease progression.

## MATERIALS AND METHODS

### Tissue collection

All liver samples were collected from healthy margins of excess surgical tissue at Massachusetts General Hospital in accordance with protocols approved by the Mass General Brigham (MGB) or Dana Farber Cancer Institute (DFCI) Institutional Review Boards (IRB) after informed consent. Samples were collected as excess tissue from surgical specimens and reviewed for histology. Samples were considered healthy if histology was normal, without inflammatory or fibrotic changes. Samples with grade 1 steatosis without inflammatory or fibrotic changes were also included in the healthy group.

### Sequencing

TST (Tween with salts and Tris) buffer was prepared as previously described [28] with final concentrations of 146mM NaCl, 10mM Tris-HCl ph7.5, 1mM CaCl2, 21mM MgCl2, 0.03% Tween-20 and 0.01% BSA in ultrapure water. From dry ice, a piece of frozen tissue was placed into a gentleMACS C Tube on dry ice (Miltenyi Biotec, cat. no. 130-093-237) with 2ml of TST buffer including 1U/ml Protector RNase inhibitor (Sigma-Aldrich, cat. no. 3335402001). Tissue was immediately dissociated using a gentleMACS Dissociator (Miltenyi Biotec, cat. no. 130-096-427) with the m_Spleen_01_01 program twice and incubated on ice for 5min to complete a 10min incubation with the TST buffer. C tubes were spun at 4 °C for 2min at 500g. The pellet was resuspended in TST buffer, filtered through a 40µm Falcon cell strainer (VWR International, LLC, cat. no. 43-50040-51) into 50ml conical tube. The strainer was washed with 1ml 1TST buffer including 0.5U/ml Protector RNase inhibitor before use. Additional 1ml of 1TST buffer including 0.5U/ml Protector RNase inhibitor was used to wash the gentleMACS C Tubes and filter. 1ml of 1XST buffer including 0.5U/ml Protector RNase inhibitor was added to the filter for a final wash. The sample was transferred to a 15ml conical tube and centrifuged at 4 °C for 10min at 500g. The pellet was resuspended in 1XPBS (-Mg/-Ca) 1% BSA, and 1U/ml Protector RNase inhibitor buffer (volume depends on pellet size, between 100-200 μl). The nucleus solution was filtered through a 35 µm Falcon cell strainer (Corning, cat. no. 352235). Nuclei were counted using a INCYTO C-chip disposable hemocytometer (VWR International, cat. no. 22-600-100). 8,000 - 12,000 nuclei of the single-nucleus suspension were loaded onto the Chromium Chips for the Chromium Single Cell 3′ Library (Chromium Next GEM Single Cell 3’ Kit v3.1, cat. no. PN-1000268, PN-1000120, PN-1000215) according to the manufacturer’s recommendations (10x Genomics). Gene expression libraries were constructed and indexed according to the manufacturer’s instructions and pooled for sequencing on either a NovaSeq 6000 or NovaSeqX sequencer (Illumina). All libraries were sequenced to a targeted depth of 400 million reads in the following configuration: R1: 28bp, R2: 90bp, I1, I2: 10bp.

### Single-nucleus RNA-seq quality control and analysis

Samples were demultiplexed using Cell Ranger mkfastq or bclconvert and aligned to the GENCODE v42 reference genome using Cell Ranger v7.0.1. Ambient RNA contamination was removed with SoupX version 1.6.2 (**Supplementary Material**), and doublets or multiplets were filtered out using DoubletFinder v2.0.4 (**Supplementary Material**). High-quality cells were integrated with Harmony v1.2.0 and clustered at different resolutions using Seurat (v4.3.0), followed by UMAP visualization (**Supplementary Material**). Clusters identified were assigned cell type labels using the ‘sctype_scor’ function from the ScType R package (**Supplementary Material**), utilizing the ‘ScTypeDB_full.xlsx’ database downloaded on 26/06/2023 (**Supplementary Material**). Marker genes were selected through differential expression analysis for each cell type. Cell-type proportions were analyzed with the speckle R package v0.99.7 using metadata for sex and age group (**Supplementary Material**).

### Differential expression, gene ontology enrichment and cell-cell communication analysis

Differential expression analysis was performed using the MAST test in Seurat v4.3.0 (FindMarkers), comparing sex (male vs. female), age groups (younger: 18-40, middle: 41-60, older: 61-90 years), and age contrasts within sex (younger vs. older females; younger vs. older males). Donor and sex (for age group comparison) or age group (for sex comparison) were included as latent variables to control for batch effects (**Supplementary Material)**. Age-related comparisons focused on pairwise analyses between younger, middle-aged, and older groups within each cell type. Genes consistently increasing or decreasing across age groups were identified as age-associated markers (**Supplementary Material)**. Genes were called significant per cell type if detected in ≥10% of cells in either group (min.pct = 0.1), showed |log2FC| ≥ 0.25, and had Bonferroni-adjusted p-values (p_val_adj) ≤ 0.1. Gene ontology enrichment for biological processes was conducted using EnrichR version 3.2 with the WikiPathway_2023_Human library (**Supplementary Material)**. CellChat v2.1.2 was used to analyze communication patterns across sex (males vs. females) and age groups (younger: 18-40 years, middle: 41-60 years, older: 61-90 years). CellChat was applied independently to each group and then merged for comparison using the mergeCellChat function (male vs. female for sex, and younger, middle, and older groups for age) (**Supplementary Material)**.

### STRING-DB network analysis

Protein-protein interactions (PPIs) were analyzed using the R package STRINGdb v2.10.1. The STRING database (species = 9606, version = 11.5) was accessed with the network type set to full (**Supplementary Material)**. Gene ontology enrichment analysis (WikiPathways) and subcellular localization analysis were conducted using STRINGdb and networks were visualized using the STRING online server (https://version-11-5.string-db.org/ (**Supplementary Material)**.

## RESULTS

### Charting the human liver landscape with single-nucleus RNA sequencing

We performed snRNA-seq to construct a map of the healthy human liver across 37 samples from female and male donors, spanning more than seven decades of life (**Table S1**). Over 195,000 nuclei were analyzed after establishing quality control metrics (**Materials and Methods**) and removing ambient RNA and doublets (**Table S1**). The dataset includes 113,271 nuclei from 20 female donors (57.9%) and 82,244 nuclei from 17 male donors (42.1%) (**Fig. 1A**). Age distributions across donors are as follows: nine samples from donors aged 18-40, containing 50,551 nuclei (25.8%) labeled as younger; 12 samples from donors aged 41–60, comprising 62,098 nuclei (31.8%) labeled as middle; and 16 samples from donors over age 60, totaling 82,866 nuclei (42.4%) labeled as older (**Fig. 1B, Fig. S1**). Nuclei from all samples were integrated, and clustering was performed at multiple resolutions (**Fig. S2**). A resolution of 0.5 with 18 clusters (0-17) was selected for downstream analysis due to its ability to capture biologically meaningful sub-clusters, as described below (**Fig. S3A, Supplementary Material**).

**Fig. 1.**
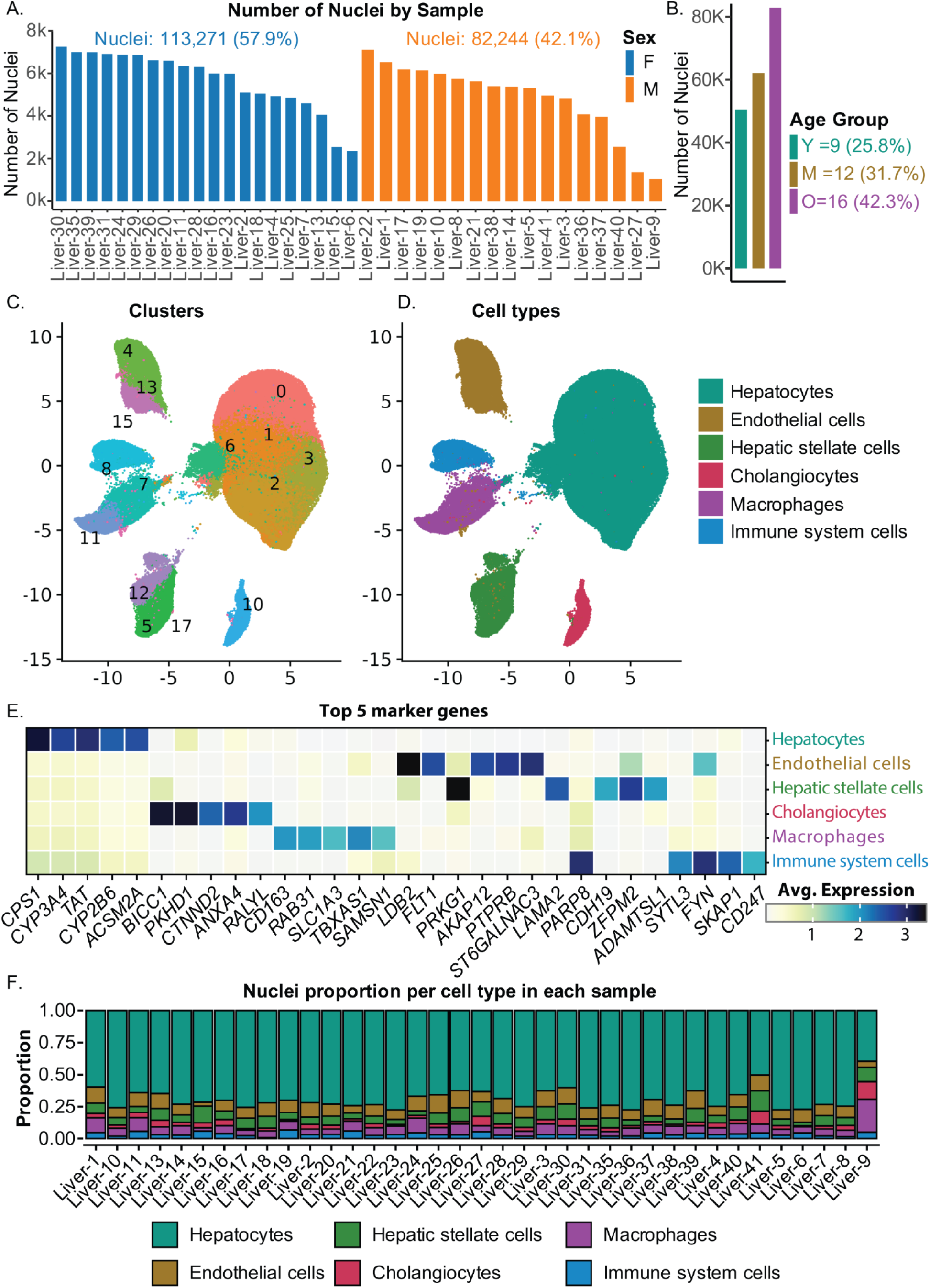
Sample composition, clustering, and cell-type proportions in the human liver. **(A)** Bars indicate nuclei counts within individual samples. Female (F) are in blue and male (M) are in orange. **(B)** Bars indicate nuclei counts by age group. Age categories are grouped as Younger (18–40 years, teal), Middle (41–60 years, mustard yellow), and Older (> 60 years, purple). **(C)** Clustering of nuclei at resolution 0.5 visualized using UMAP. Each point represents an individual nucleus, colored according to its assigned cluster. Cluster identities are labeled numerically to correspond with specific populations identified in the analysis. **(D)** Cell-type assignments for each cluster (from C), identified using ScType, are displayed on the UMAP plot. **(E)** Top 5 marker genes defining each cell type identified in panel D are shown. **(F)** This panel illustrates the proportions of each cell type across all samples.

As part of the cluster quality control process, we quantified *MALAT1* expression (**Table S2, Fig. S3B**). *MALAT1* is a long noncoding RNA retained in the nucleus, allowing it to serve as an indicator of high-quality nuclei [29,30]. We excluded two clusters (9 and 14) from downstream analysis because there was enrichment of nuclei with low *MALAT1* expression (fewer than three reads per nucleus). We also excluded cluster 16 from downstream analyses, as all 167 cells originated from a single donor sample (sample 15; **Table S2)**. Fifteen clusters remained in the final UMAP (**Fig. 1C)**.

Clusters were initially annotated using ScType [31] with liver as the target tissue, and assigning clusters to hepatocytes, endothelial cells, hepatic stellate cells (HSCs), cholangiocytes, macrophages, and non-macrophage immune cell populations (**Fig. 1D**, **Table S3**). We further validated our annotations using CellTypist [32] (**Fig. S4, Table S3**). HSCs were not included in the CellTypist annotation, and this cluster was labeled as fibroblasts. We retained the HSCs label, as this is more specific to the liver annotation. CellTypist contained more detailed annotation of immune cells than the liver annotation for ScType, and the cluster labeled as non-macrophage immune cells by ScType was labeled as T cells by CellTypist (**Supplementary Material**).

Marker genes for each annotated cell type were determined using the FindAllMarkers function, showcasing the unique gene expression profiles that characterize these populations (**Fig. 1E, Table S4**). The top five genes enriched in hepatocytes include cytochrome p450 family members *CYP3A4* and *CYP2B6,* along with *CPS1* and *TAT*. Endothelial cells are enriched in expression of genes that include the phospho-signaling molecules *FLT1, AKAP12,* and *PTPRB*, while genes most enriched in HSCs include *LAMA2, ADAMTSL1, and CDH19*, which are involved in the extracellular matrix or cell adhesion. For cholangiocytes, the most enriched genes include adhesion molecules *CTNND2* and *PKDH1* along with *ANXA4,* which is linked to exocytosis. Immune cells were separated into macrophages and non-macrophage populations. Macrophages were enriched in expression of the scavenger receptor *CD163*, the phagocytic factor *RAB31*, and *TBXAS1*, responsible for production of thromboxane *A2*. Non-macrophage immune cells as a group show increased expression of genes including *SKAP1* and *CD247 and FYN*, which encode products involved in lymphocyte receptor signaling. Each cell type was represented in each sample, ensuring that downstream analyses included representation from every sample (**Fig. 1F, Table S3**).

### Evaluating hepatocyte zonation

We next classified hepatocytes into portal, central, and mid-zone based on the expression of key zonal markers: *ASS1*, *ASL*, *GLUL* and *CYP2E1 [33]*. We calculated the Central-to-Portal Ratio (CPR) by subtracting the summed expression of the markers *ASS1* and *ASL* (portal; zone 1) from the summed expression of *GLUL* and *CYP2E1* (central; zone 3) for each hepatocyte sub-cluster (**Fig. 1C**). This ratio provides a quantitative measure of the zonal distribution. Clusters were categorized as portal if their CPR values were less than or equal to the lower quantile (clusters 2, 3), central if their CPR values were greater than or equal to the upper quantile (clusters 0, 6), and mid-zone for in between values (cluster 1) (**Fig. 2A, Table S5**). This classification was validated with UMAP visualization and confirmed with density plots, demonstrating distinct zonal distributions (**Fig. 2B**, **2C**). Portal, mid-zone, and central hepatocytes were then annotated within the overall UMAP (**Fig. 2D**).

**Fig. 2.**
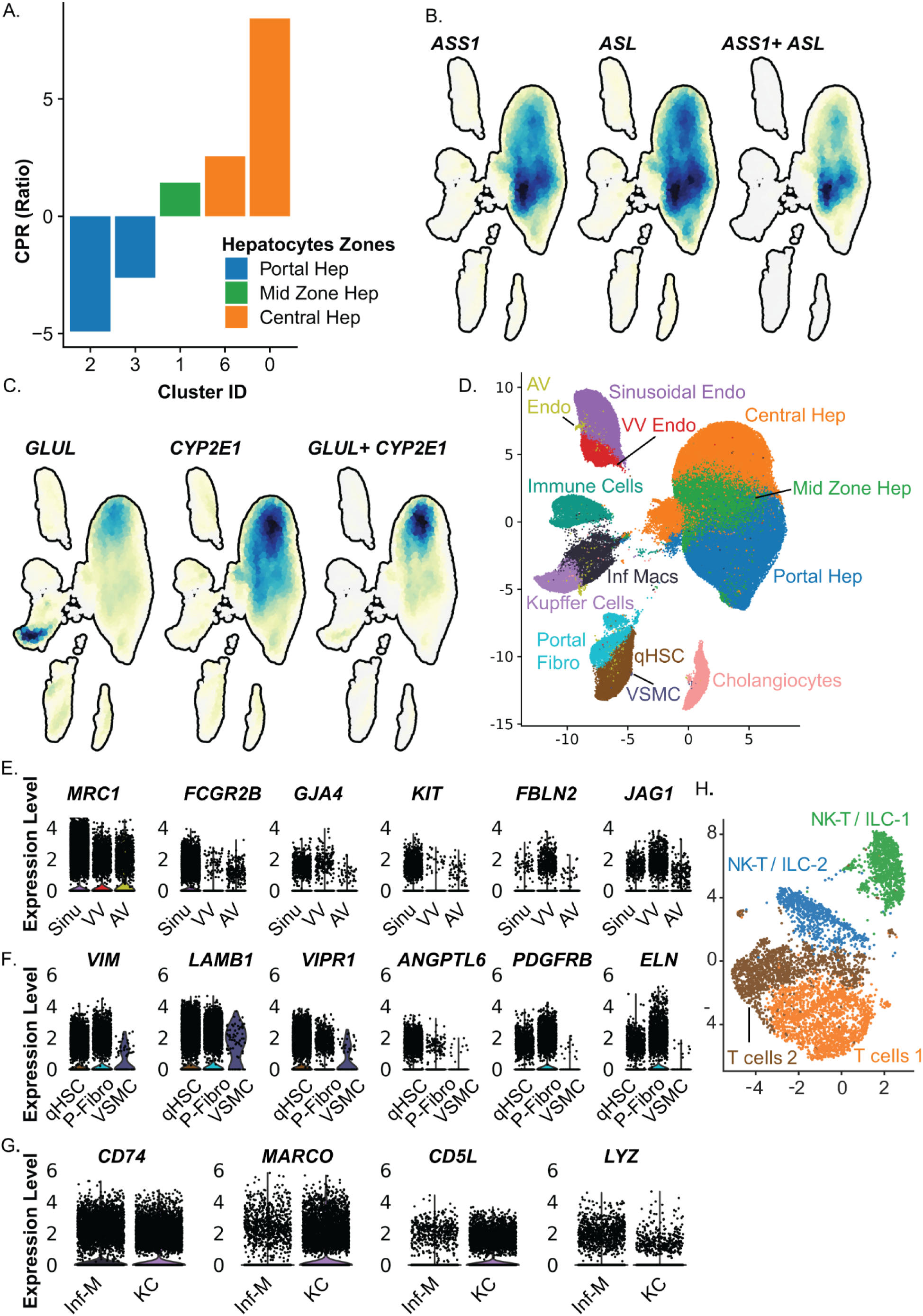
Defining hepatocyte zones and subtypes in non-hepatocytes. **(A)** Central-to-Portal Ratio (CPR) for hepatocyte clusters, calculated using zonal markers *ASS1*, *ASL* (portal zone), and *GLUL*, *CYP2E1* (central zone). Clusters were classified as portal (CPR values ≤ lower quantile: clusters 2, 3), central (CPR values ≥ upper quantile: clusters 0, 6), or mid-zone (intermediate CPR values: cluster 1). **(B)** Density plots showing expression of *ASS1* and *ASL* overlaid on the UMAP, alongside a combined (joint) density plot for their co-expression. **(C)** Density plots showing expression of *GLUL* and *CYP2E1* overlaid on the UMAP, along with a combined (joint) density plot reflecting their co-expression distribution. **(D)** UMAP visualization of cell types and subtypes, detailing hepatocytes, non-parenchymal cells, and immune populations. **(E-G)** Marker genes identifying subtypes of indicated populations as shown in panel D. **(H)** UMAP visualization of non-macrophage immune subtypes.

### Defining non-hepatocytes cell types and subtypes

We then evaluated each individual non-hepatocyte cluster (**Fig. 1C**) to define cell types and subtypes. We identified sinusoidal, venous vascular (VV), and arterial vascular (AV) endothelial cells [34] (**Fig. 2E, Table S6**). HSCs were separated into quiescent HSCs (qHSCs) and portal fibroblasts [35], and a population of vascular smooth muscle cells (VSMCs) was annotated (**Fig. 2F**). Macrophages were separated into Kupffer-like cells (*CD74^high^*, *MARCO^high^*, *CD5L^high^*) and inflammatory macrophages (*CD74^high^*, *LYZ^high^*, *MARCO^low^*) [1] (**Fig. 2G**). Cell types were evenly distributed across samples (**Fig. S5**), with balanced representation across sexes (**Fig. S6**) and age groups (Younger, Middle, and Older) **(Fig. S7**). These annotations highlight the cellular heterogeneity within these populations captured across sexes and over many decades of age. Sample 9 did show an increase in the fraction of inflammatory macrophages (**Fig. S5**). Because this sample contained the smallest number of nuclei (**Fig. 1A**), it contributed only 2.86% to the total inflammatory macrophage population but still included the other abundant cell types, so it was retained for analysis.

The non-macrophage immune system cell population identified (6,544 cells) was then sub-clustered from the raw data using the same analysis pipeline and annotated with the ScType tool using Immune System Cells as the target tissue to define subtypes of non-macrophage immune cells. This approach identified two populations of T cells and two populations of NK T-like cells. (**Fig. 2H, Table S7**). We further validated these cell type annotations with CellTypist [32] (**Fig. S8**, **Table S7**). The Celltypist analysis confirmed the two T cell populations. Celltypist does not contain annotations for NK T-like cells and instead annotated these clusters as innate lymphoid cells (ILCs), which include NK cells (**Supplementary Material**). To acknowledge the two different annotations for these clusters, they are labeled as NK-T / ILC populations. This sub-clustering provides greater resolution of the non-macrophage immune cell landscape within the liver.

### Effect of sex on differential gene expression and biological pathways by cell type and subtype

Cell type proportions were evaluated [36] (**Materials and Methods**) between females (20) and males (17), and no significant differences were identified with the exception of AV endothelial cells, which were increased in females compared to males (0.71% vs 0.45%; FDR = 0.077), supporting differential expression analysis to uncover sex-specific gene expression differences (**Table S8**). Due to low cell counts, AV endothelial cells (830 female cells vs 400 male cells) and VSMCs (49 female cells vs 27 male cells) were not included in the overall analysis. Data for AV endothelial cells are contained in **Table S9**, and there were no differentially expressed genes identified for VSMCs. Non-macrophage immune cells were analyzed collectively because of the smaller size of individual subgroups (**Table S8**).

Differential expression analysis across cell types and subtypes was performed with donor and age as co-variates and revealed 425 genes upregulated in females and 581 genes upregulated in males, with 37 genes demonstrating upregulation in both sexes but in different cell types or subtypes (**Table S9**). These findings suggest that while most genes that change in expression are either enriched in female cell types or male cell types, a smaller fraction of genes can be increased in different cell types depending on whether the cells are female or male. When focusing exclusively on autosomal genes (excluding X and Y chromosomes), 374 genes were upregulated in females, 520 in males, and 36 genes were upregulated across both sexes, again in distinct cell types. We then focused on autosomal genes for the subsequent analyses. The top four most enriched genes in female and male cell types and subtypes were plotted (**Fig. 3A**), highlighting genes that are enriched in specific cell types and subtypes as well as those shared across multiple categories.

**Fig. 3.**
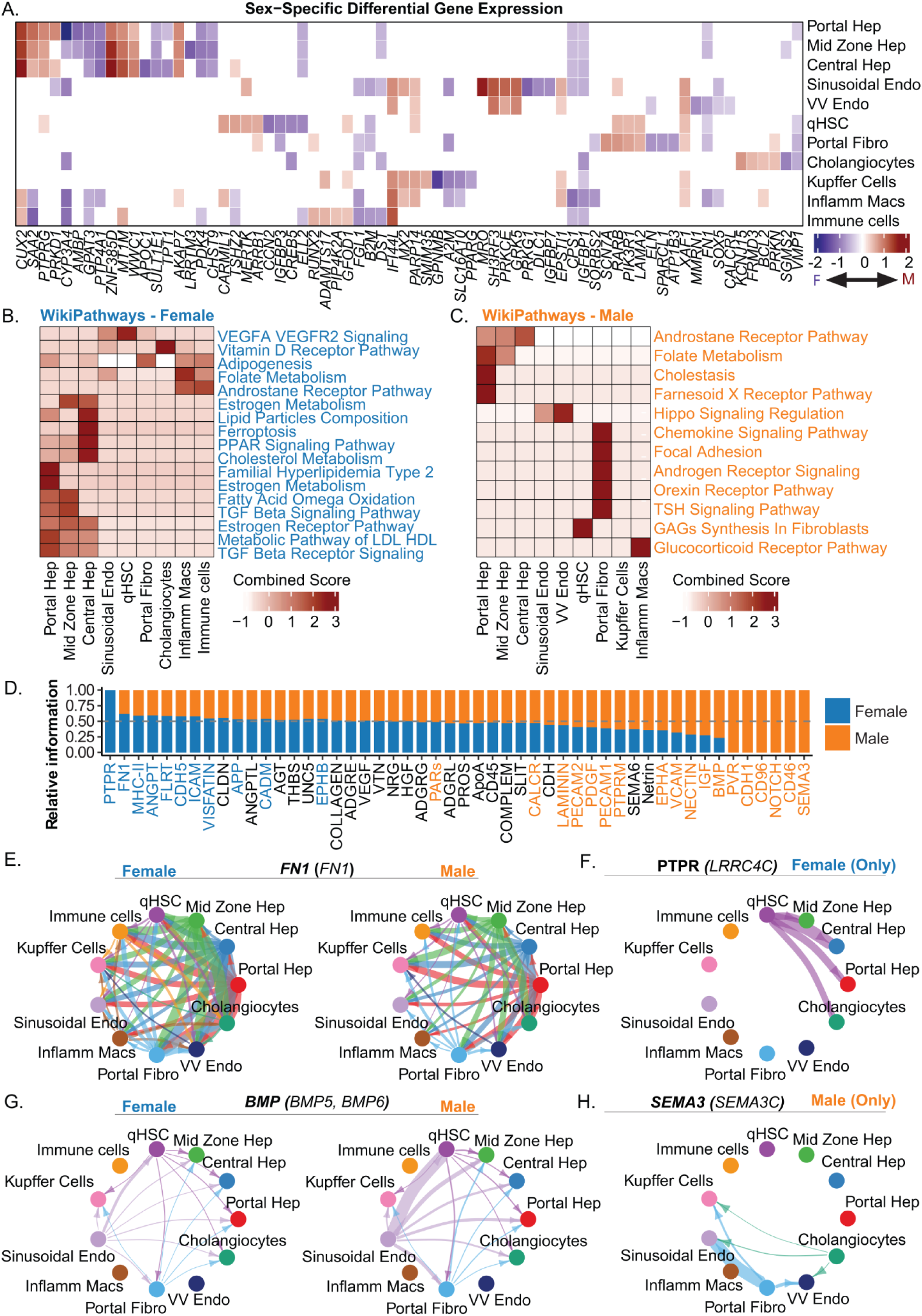
Sex-specific gene expression, pathway enrichment, and signaling. **(A)** Heatmap displaying the log2 fold change of gene expression between females and males across cell types and subtypes within the liver. Red indicates higher expression levels in males, while blue signifies higher expression levels in females. The top 4 differentially expressed genes, upregulated in each group (female or male) and expressed in at least 25% of cells are displayed for each cell type and subtype. Genes were called significant in each cell type if detected in ≥10% of cells in either group (min.pct = 0.1), showed |log2FC| ≥ 0.25, and had Bonferroni-adjusted p-values (p_val_adj) ≤ 0.1. **(B)** Biological processes (Wiki pathways) enriched in differentially expressed genes across cell types in females. The combined score was calculated by logarithmically transforming the p-value from Fisher’s exact test and multiplying it by the z-score. Enriched functional categories were considered significant when the false discovery rate (FDR) was ≤ 0.1 and each term included at least three contributing genes. **(C)** Biological processes (Wiki pathways) enriched in differentially expressed genes across cell types in males. Glycosaminoglycans is abbreviated as GAGs. Enriched functional categories were considered significant when the false discovery rate (FDR) was ≤ 0.1 and each term included at least three contributing genes. **(D)** Significant signaling pathways (p ≤ 0.05) were ranked based on differences in overall information flow within the inferred networks between females and males. The top signaling pathways labeled in blue are more enriched in females. Pathways labeled in orange are more enriched in males. Pathways labeled in black are not statistically enriched in either category. **(E)** Chord circle plots display significantly interacting pathways and communication probabilities of FN1 signaling (higher in females compared to males), with enriched ligand (FN1). **(F)** Chord circle plot displays significantly interacting pathways and communication probabilities of PTPR signaling, (present only in females), with enriched ligand (LRRC4C). **(G)** Chord circle plot displays significantly interacting pathways and communication probabilities of BMP signaling(higher in males as compared to females), with enriched ligands (BMP5 and BMP6). **(H)** Chord circle plot displays significantly interacting pathways and communication probabilities of SEMA3 signaling (present only in males), with enriched ligand (SEMA3C).

We next examined the biological processes enriched in cell types based on differential expression between female and male cells (**Fig. 3B** and **Table S10**). The Estrogen Receptor Pathway (WP2881) and Estrogen Metabolism (WP697, WP5276) were enriched in female hepatocytes. TGF-beta Receptor Signaling (WP560) was enriched in all hepatocytes, with the strongest enrichment in portal hepatocytes, and the TGF-beta Signaling Pathway (WP366) was enriched in portal and mid-zone hepatocytes. Multiple pathways related to lipid metabolism were increased in female cells. Lipid Particle Composition (WP3601) and Cholesterol Metabolism (WP5304) showed the strongest enrichment in central hepatocytes, while Metabolic Pathway of LDL, HDL and TG (WP4522) was enriched across all hepatocytes. Fatty Acid Omega Oxidation (WP206) was enriched in portal and mid-zone hepatocytes, and Familial Hyperlipidemia type 2 (WP5109) was enriched in portal hepatocytes. Vitamin D Receptor Signaling (WP2877) was enriched in cholangiocytes, VEGFA-VEGFR2 Signaling (WP3888) in qHSCs, and Folate Metabolism (WP176) in inflammatory macrophages and non-macrophage immune cell populations.

In male cells, the Farnesoid X Receptor Pathway (WP2879) was enriched in portal hepatocytes, while Chemokine Signaling (WP3929), the Orexin Receptor Pathway (WP5094), and Focal Adhesion (WP306) were enriched in portal fibroblasts. The Glucocorticoid Receptor Pathway (WP2880) was enriched in inflammatory macrophages, whereas the Hippo Signaling Regulation Pathway (WP4540) was enriched in both sinusoidal and VV endothelial cells (**Fig 3C** and **Table S10**).

There were also examples of common pathways enriched in different cell types or subtypes between females and males, including the Androstane Receptor Pathway (WP2875), which is most enriched in inflammatory macrophages and (non-macrophage) immune cells in females, while enriched in hepatocytes in males (**Fig. 3B**, **3C**). Folate Metabolism (WP176) is also enriched in inflammatory macrophages and (non-macrophage) immune cells populations in females but portal and midzone hepatocytes in males (**Fig. 3B**, **3C**), while Focal Adhesion PI3K Akt mTOR Signaling (WP3932) was enriched in female inflammatory macrophages and male portal fibroblasts (**Table S10**). These findings highlight both female- and male-specific pathways enriched within cell types in the healthy liver and how even the same pathways may modulate the activity of different cell types between female and male individuals.

### Impact of sex on cell-cell communication

Given the differences in gene expression and pathways enriched between female and male cells, we next predicted signaling pathway strengths by assessing communication probabilities among cell groups using CellChat (v2). This revealed distinct pathways enriched in female and male cells, as well as shared pathways (**Fig. 3D**, **Table S11**). Female cells showed enrichment in signaling networks that include PTPR, ANGPT, FN1, and VISFATIN (**Fig. 3D**, blue labels), while male cells were enriched in pathways including SEMA3, VECAM, IGF, BMP, and PDGF (orange labels). Additional pathways are active but showed no significant sex-specific differences (**Fig. 3D**, black labels). These findings suggest sex-based gene expression differences can influence cellular responses and outputs to identical stimuli.

We next visualized cell-cell communication across pathways in female and male cells (**Table S11**). In females, FN1 signaling was stronger with more extensive connections compared to males (**Fig. 3E**). FN1 signaling from hepatocytes was enriched in females, but the majority of FN1 produced by hepatocytes is secreted into the plasma [37], so the impact on other resident liver cell types is not clear. We also visualized cell-cell communications without hepatic FN1 (**Fig. S9**), which showed that FN1 signaling remained enriched in females, with portal fibroblasts signaling to VV and sinusoidal endothelial cells, and immune cell populations in addition to FN1 signaling from cholangiocytes, immune cells, and inflammatory macrophages. PTPR signaling is only enriched between female cells, with qHSCs signaling to hepatocytes and cholangiocytes in females but not males (**Fig. 3F**). In contrast, BMP signaling was enriched in males, with the strongest connection from sinusoidal endothelial cells to qHSCs (**Fig. 3G)**. The SEMA3 pathway was more active in males as well, signaling primarily from portal fibroblasts and cholangiocytes to VV and sinusoidal endothelial cells as well as Kupffer cells, without active signals identified in females (**Fig. 3H**). These results show that signaling pathways can operate at different strengths and involve different cell types within the female and male liver.

To identify molecules driving cell-cell interactions, we next visualized ligand and receptor (L-R) pairs enriched in female and male cell types (**Fig. 4A-D, Table S11**). Pairs were excluded if the receptor-expressing cell type showed higher ligand expression than the ligand-expressing cell type, because we could not exclude that the primary signal was autocrine. We also removed L-R pairs that were enriched in both female and male livers but involved different cell types to simplify visualization. The analysis focused on signaling from portal hepatocytes and sinusoidal endothelial cells as examples, and the full results are in **Table S11**.

**Fig. 4.**
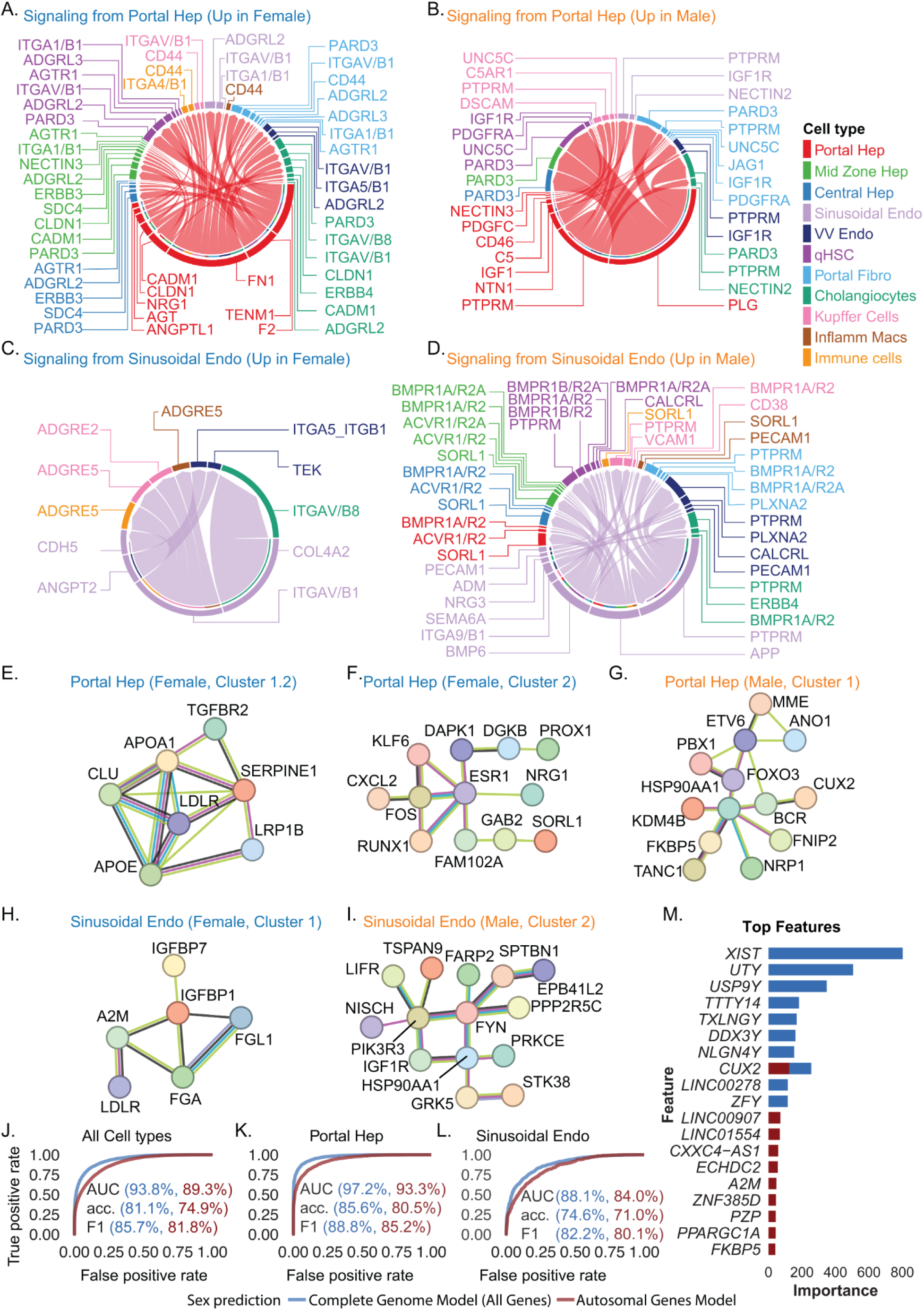
Sex-specific intercellular interactions, protein networks, and application of machine learning. **(A-D)** Chord diagrams representing sex-specific ligand-receptor (L-R) interactions. Outer rings denote different cell types, while links illustrate interactions, with ligands expressed by the source cell type and receptors expressed by the target cell type. L-R signaling originating from portal hepatocytes enriched in females (**A**), and males (**B**). L-R signaling originating from sinusoidal endothelial cells enriched in females (**C**) and males (**D**). **(E-H)** STRINGDB network analysis of protein-protein interactions (PPIs) Nodes represent proteins, while edges (lines) depict interactions, with different colors indicating evidence from distinct sources (Red: fusion evidence, Green: neighborhood evidence, Blue: co-occurrence evidence, Purple: experimental evidence, Yellow: text mining evidence, Light blue: database evidence, Black: co-expression evidence). **(E)** Protein network in female portal hepatocytes enriched in pathways related to lipid metabolism. **(F)** Protein network in female portal hepatocytes enriched in estrogen signaling and oncostatin M signaling pathways. **(G)** Protein network in male portal hepatocytes. **(H)** Protein network in female sinusoidal endothelial cells. **(I)** Protein network in male sinusoidal endothelial cells. **(J-L)** Machine learning-based prediction of sex using **(J)** all cell types combined, **(K)** portal hepatocytes, and **(L)** sinusoidal endothelial cells. The blue line indicates the AUC for models trained on all genes (including sex chromosomes), while the red line represents models trained exclusively on autosomal genes. **(M)** The plot shows the top 10 features for predicting sex in the Complete Genome Model (blue bars) and the Autosomal Gene Model (red bars). The y-axis represents feature importance scores, with higher values indicating greater contributions.

In females, enriched L-R pairs include the combination of ANGPTL1 (ligand) expressed by portal hepatocytes and ITGA1_ITGB1 (receptor complex) expressed by sinusoidal endothelial cells, qHSCs, portal fibroblasts, and mid-zone hepatocytes (**Fig. 4A**). TENM1 (ligand) also paired with ADGRL3 (receptor) in qHSCs and portal fibroblasts, and CADM1 (ligand) paired with NECTIN3 (receptor) in mid-zone hepatocytes. FN1 (ligand) from portal hepatocytes also displayed potential interactions across multiple cell types through integrin receptor complexes (ITGA4_ITGB1, ITGA5_ITGB1, and ITGAV_ITGB1), CD44, and SDC4.

In contrast, IGF1 (ligand) from portal hepatocytes pairs with IGF1R (receptor) on sinusoidal and VV endothelial cells in males (**Fig. 4B**). PLG (ligand) pairs with PARD3 (receptor) in qHSCs, portal fibroblasts, cholangiocytes, and hepatocytes, while NTN1 (ligand) from portal hepatocytes interacts with DSCAM (receptor) on Kupffer cells and UNC5C (receptor) on qHSCs, portal fibroblasts, and Kupffer cells. These findings highlight the diversity of signaling involving hepatocytes between females and males including pathways in cell adhesion in females (TENM1, CADM1) and repair/regeneration (PLG and NTN1) in males.

Enrichment of L-R pairs in females involving signaling from sinusoidal endothelial cells include ANGPT2 (ligand) combined with TEK and ITGA5_ITGB1 (receptors) expressed by VV endothelial cells and COL4A2 (ligand) paired with ITGAV_ITGB8 (receptor complex) expressed by cholangiocytes (**Fig. 4C**). In males, enriched L-R pairs include BMP6 (ligand) expressed by sinusoidal endothelial cells paired with ACVR1_ACVR2A (receptor complex) in mid-zone hepatocytes, ACVR1_ACVR2A/BMPR2 in portal and central hepatocytes, and BMPR1A_ACVR2A in mid-zone hepatocytes, qHSCs, and portal fibroblasts (**Fig. 4D**). In addition, the combination of APP (ligand) expressed by sinusoidal endothelial cells and SORL1 (receptor) expressed by inflammatory macrophages, hepatocytes, and non-macrophage immune cells is also enriched in males. Taken together, the L-R analyses show how individual L-R pairs can differ between liver cell types and subtypes in the female and male liver, establishing different circuitry to respond to both normal physiological signals and those associated with injury and disease.

### Influence of sex on protein networks

We next predicted differences in protein-protein interactions between female and male cells in the liver based on gene expression data through STRINGDB network analysis. We focused on networks within hepatocytes and endothelial cells for visualization and included all data for interactions and subcellular compartments in **Table S12**. In female portal hepatocytes, we identified a sub-network containing APOA1, APOE, LDLR, and LRP1B (**Fig. 4E**); this network was enriched in pathways involved in Lipid Particle Composition (WP3601), Metabolic Pathways of LDL, HDL, and TG (WP4522), and Familial Hyperlipidemia-related Pathways (WP5108, WP5109, WP5110, WP5111, WP5112) (**Fig. 4E**). Subcellular localization analysis further revealed that the associated proteins are primarily localized to lipoprotein particles (GOCC:0034362, GOCC:0062136) or are lipoprotein receptors (LDLR and LRP1). A protein network in female portal hepatocytes was also identified that centered on Estrogen Receptor 1 (ESR1) and FOS (**Fig. 4F**). This network was enriched in Estrogen Signaling (WP712), and ESR1-containing networks were also present in central and mid-zone hepatocytes (**Fig. S10**). Proteins in this network are primarily localized to the nucleus and include nuclear receptors and intermediate signaling molecules.

An additional protein network enriched in female portal hepatocytes included ADH1A, ADH1B, CYP3A4, SULT1E1, and GSTA1 (**Fig. S9)** and contained components of Fatty Acid Omega Oxidation (WP206) and Estrogen Metabolism (WP697). Proteins in this network are primarily cytoplasmic (GOCC:0005737). Male portal hepatocytes displayed a network including HSP90AA1 and FKBP5 (**Fig. 4G**), which modulate nuclear receptor signaling [38,39], and the proteins identified in this network are also primarily cytoplasmic.

Female sinusoidal endothelial cells showed enrichment of a protein network containing FGA, FGB and FGG, proteins linked to Folate Metabolism (WP176), the Blood Clotting Cascade (WP272), and Fibrin Complement Receptor 3 Signaling (WP4136) (**Fig. 4H**). Subcellular localization analysis places this network within the extracellular space (GOCC:0005615). A network containing PIK3R3, IGF1R, HSP90AA1,and PPP2R5C, which are all members of the PI3K/Akt Signaling Pathway (WP4172), was enriched in male sinusoidal endothelial cells (**Fig. 4I**), and subcellular localization analysis suggests these interactions occur primarily in the cytosol (GOCC:0005829). Together, these results highlight differences in protein-protein interaction networks between female and male cell types that can affect their intrinsic activity and response to external signals.

### Applying machine learning to evaluate male and female cell populations

To determine if liver cells can be classified as female or male by machine learning, we applied a Random Forest classifier across all cell types using 80% of the data for training and 20% for testing (**Supplementary Material**). Two models were trained: one using all genes as a baseline (full model) and another excluding X and Y chromosome genes (autosomal model). The full model achieved an accuracy of 81.1% and recall of 97.9%, while the autosomal model still performed well, with 74.9% accuracy and 97.3% recall (**Fig. 4J**). We then evaluated classification performance for individual cell types and subtypes. Both models performed robustly, with the full model achieving slightly higher accuracy and recall. For example, portal hepatocytes were classified with 85.6% accuracy and 99.4% recall in the full model and 80.5% accuracy and 98.0% recall in the autosomal model (**Fig. 4K**). Sinusoidal endothelial cells showed similar trends, with 74.6% accuracy and 97.4% recall in the full model and 71.0% accuracy and 96.7% recall in the autosomal model (**Fig. 4L**). Other cell types also demonstrated robust classification performance (**Fig. S12A-I**). The autosomal model achieved similar accuracy to the full gene model across all cell types, demonstrating that cell type and subtype classification can be effectively performed using only autosomal gene expression (**Table S13** and **Fig. S12A-I**).

These results highlight the key role of autosomal genes in differentiating female and male cells. When the full model was run using all cell types, X- and Y-linked genes such as *XIST*, *UTY*, *TTTY14*, and *USP9Y* were among the top features, as expected. *XIST*, known for its critical role in X-chromosome inactivation in females [40], also exhibited the highest feature importance score. However, even after excluding X and Y chromosome genes, the classifier highlighted key genes contributing sex differences within the liver, including *CUX2* (**Fig. 4M**).

Our analysis identified increased expression of *CUX2* in males, while murine models detect increased *CUX2* in female livers [41]. To validate our findings, we analyzed *CUX2* expression using bulk RNA-seq data from the GTEx Portal (v8) [42]. Across 14 GTEx tissues where *CUX2* expression was detectable (median transcripts per million (TPM) > 1), we also observed male-biased expression in the liver and regions of the brain (**Fig. S13** and **Table S14**). In the human liver, the median TPM was 13.8 in males vs. 7.5 in females, confirming a male bias in bulk samples. Similarly, in cortical brain regions, male medians exceeded female values (e.g., Brain Cortex: 16.4 vs. 15.5; Frontal Cortex BA9: 15.4 vs. 11.8). A few tissues demonstrated female-biased medians, including the amygdala (2.05 female vs. 1.72 male) and substantia nigra (1.91 female vs. 1.38 male). These findings further support the identification of *CUX2* as enriched in liver cell types in male donors, and reinforces the potential role of autosomal genes in regulating sex-specific functions within liver cells.

To evaluate whether the accuracy of sex prediction was influenced by donor age, we repeated the analysis using only autosomal genes within three age groups: younger donors (18-40 years), middle-aged donors (41-60 years), and older donors (>60 years) (**Supplementary Material**). Nuclei sex was predicted with 86.3% accuracy and 100% recall in younger donors, 82.9% accuracy and 98.4% recall in middle-aged donors, and 87.0% accuracy and 93.4% recall in older donors (**Table S15**).

### Effect of age on differential gene expression and biological pathways by cell type and subtype

We next investigated how gene expression changes with age across cell types and subtypes (**Fig. 1B, Fig. S1**). Age groups (younger, middle, and older) showed similar representation of cell types (**Fig. S7**), and we performed differential expression analysis, using donor and sex as co-variates, to define age-specific changes in gene expression (**Table S16**). We identified 478 genes that increase in expression from younger to middle-age to older, and 50 genes that decrease with age when analyzed by cell types and subtypes. The top five most enriched genes in older (red) versus younger cells (blue) for each cell type are shown (**Fig. 5A**). Genes enriched with age include *OAT*, *PEX6*, *BHMT*, and *PDE3B* in hepatocytes, and *PDK4* in both hepatocytes and qHSCs; the products of these genes regulate amino acid, very long-chain fatty acid, homocysteine, glucose, and glucose/cholesterol metabolism, respectively [43–47]. Sinusoidal endothelial cells, portal fibroblasts, cholangiocytes, and Kupffer cells all show differential expression of unique genes with changes in age (**Fig. 5A**, **Table S16**).

**Fig. 5.**
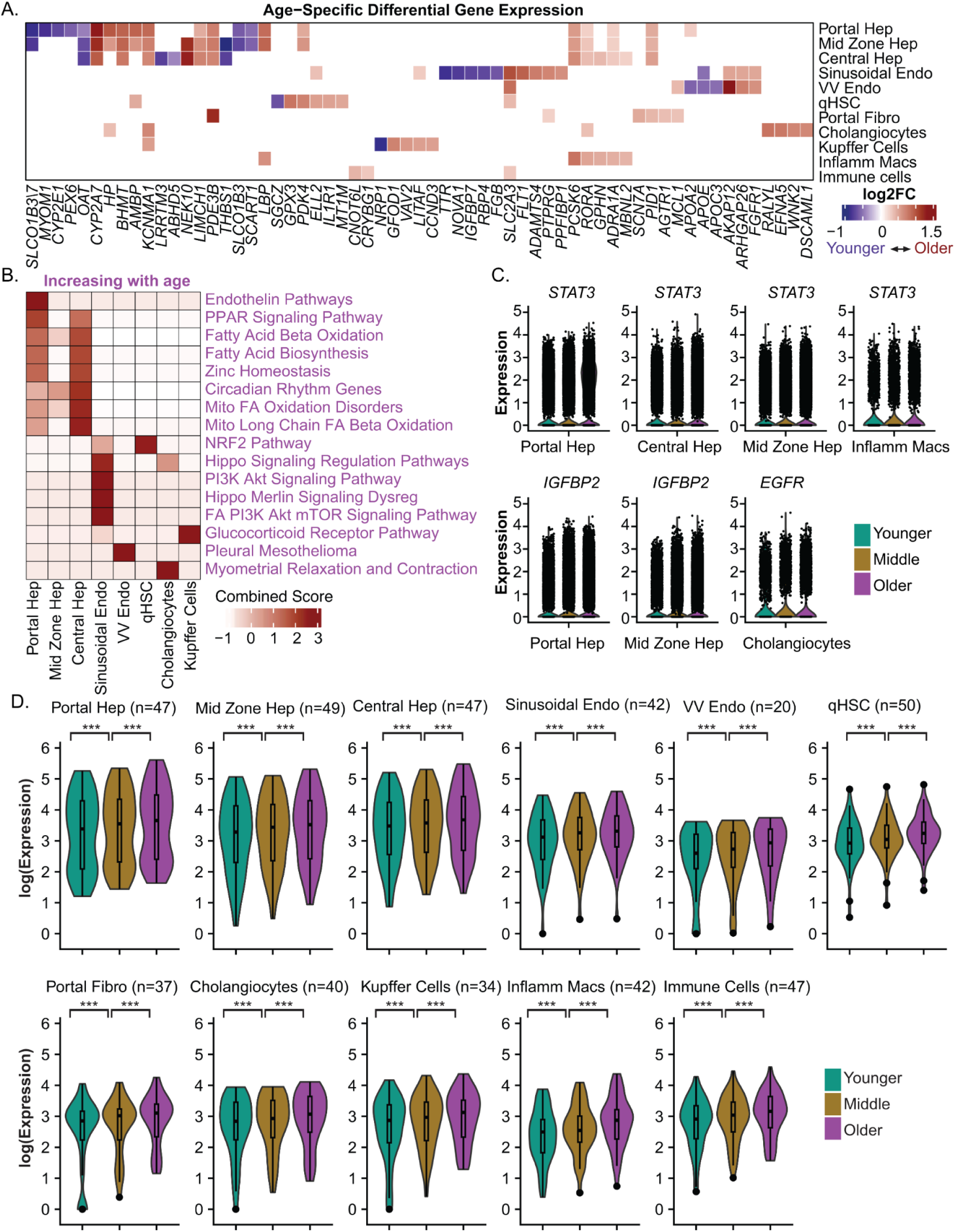
Age-related gene expression changes and senescence signatures. **(A)** Heatmap displaying the log2 fold change of gene expression between age groups across cell types. Red indicates increase in expression with age and blue decrease in expression with age. A maximum of 5 differentially expressed genes, upregulated in each group (increasing or decreasing with age) and expressed in at least 25% of cells are displayed for each cell type and subtype. Genes were called significant in each cell type if detected in ≥10% of cells in either group (min.pct = 0.1), showed |log2FC| ≥ 0.25, and had Bonferroni-adjusted p-values (p_val_adj) ≤ 0.1. **(B)** Heatmap presents a gene ontology enrichment analysis of up-regulated genes across different age groups, focusing on pathways that increase with age. Enrichr combined score is calculated by the logarithmic transformation of the p-value obtained from Fisher’s exact test, multiplied by the z-score. Enriched functional categories were considered significant when the false discovery rate (FDR) was ≤ 0.1 and each term included at least three contributing genes. **(C)** Violin plots in each panel show gene expression across three age groups: Younger, Middle, and Older. These plots highlight statistically significant changes in senescence-associated genes[49] identified in the dataset. Genes were considered significant within cell type if they were detected in ≥10% of cells in either the Younger or Older group (min.pct = 0.1), showed an absolute log2fold change (|log2FC|) ≥ 0.25, and had Bonferroni-adjusted p-values (p_val_adj) ≤ 0.1 when comparing Younger and Older groups. In addition, expression in the Middle group was required to fall between the levels observed in the Younger and Older groups. **(D)** Cell type-specific senescence signatures. Violin plots of senescence gene expression across indicated groups, with average expression values displayed on a log scale. *** p < 0.001 as obtained from pairwise Wilcoxon tests (**Materials and Methods**).

Pathways enriched with increasing age were also identified (**Fig. 5B)**. Portal hepatocytes showed enrichment for Endothelin Pathways (WP2197). Portal and central hepatocytes were enriched for Fatty Acid Biosynthesis (WP357), Mitochondrial Fatty Acid Oxidation Disorders (WP5123), and Mitochondrial Long Chain Fatty Acid Beta Oxidation (WP368), while central and mid-zone hepatocytes were enriched for Circadian Rhythm Genes (WP3594), and all hepatocytes were enriched for Fatty Acid Beta Oxidation (WP143). Sinusoidal endothelial cells were enriched with age in pathways including Hippo Signaling (WP4540, WP4541) and PI3K-Akt Signaling (WP3932, WP4172). qHSCs showed significant enrichment for the NRF2 Pathway (WP2884), and the Glucocorticoid Receptor Pathway (WP2880) was enriched in Kupffer cells. Fewer pathways were identified as enriched at a younger age due to the smaller number of genes in this group, but Lipid Particle Composition (WP3601) and Cholesterol Metabolism (WP5304) pathways were enriched in VV endothelial cells in younger donors. Collectively, this analysis underscores the cell-type-specific adaptations across key metabolic and signaling pathways with increasing age (**Table S17)**.

While data on menopause were not available for older female donors, we attempted to gain insight into possible effects of menopause by comparing female donors 40 years of age and younger with donors over 60 years of age (**Table S18)**. These ages were selected because the mean age of menopause is 51.4 years with <10% of women reaching menopause after age 56 and <2.5% reaching menopause before the age of 45 [48]. Younger donors showed enrichment in Estrogen Metabolism Signaling (WP712, WP697, WP691) and Lipid/PI3K–AKT–mTOR programs (WP5368, WP4723, WP4724), whereas older donors showed enrichment in Fatty Acid Beta-oxidation (WP143), Cholesterol Biosynthesis (WP197), and Glycolysis (WP2456) **(Table S19**). A parallel analysis in males revealed similar age-related shifts, including enrichment of Fatty Acid Beta-oxidation (WP143), Cholesterol Production (WP5304), Familial Hyperlipidemia Type 2 (WP5109), and PPAR signaling (WP3942) increased in older males. These pathways shared between aging female and male liver cells are consistent with aging, rather than menopause, as the primary driver (**Table S20 and TableS21**).

### Senescence signatures with increasing age

Expression of senescence genes increases with age in many organs, but we did not identify a senescence signature in our pathway analysis (**Fig. 5B**). We next evaluated expression of genes associated with senescence [49] to better understand how these genes change with increasing age across cell types and subtypes. We found that *STAT3* was enriched with age in all hepatocytes and inflammatory macrophages (**Fig. 5C**), consistent with induction of *STAT3* in cellular senescence in liver fibrosis and in senescent macrophages [50,51]. Genes induced with age also included *IGFBP2* in portal and mid-zone hepatocytes, as well as *EGFR* in cholangiocytes (**Fig. 5C, S14**). While only seven senescence-associated genes were significantly enriched with age within individual cell types or subtypes, different combinations of senescence genes followed the trend of increasing expression from younger to middle to older age groups for each cell type and subtype (**Table S22** and **Supplementary Material**). Using the set of genes that increase with age within each cell type and subtype, we then defined cell-type-specific senescence signatures (**Fig. 5D, Table S22**).

### Impact of age on cell-cell interactions

We next evaluated signaling pathway strength from younger to older liver cells using CellChat. In younger cells, pathways involving IGF, THBS, and CHEMERIN were enriched (**Fig. 6A**, green labels), while older cells showed enrichment in pathways related to ICAM, SPP1, SEMA6, and NRG (purple labels). Additional pathways (black labels) were active but did not significantly differ between younger and older cells (**Table S23**).

**Fig. 6.**
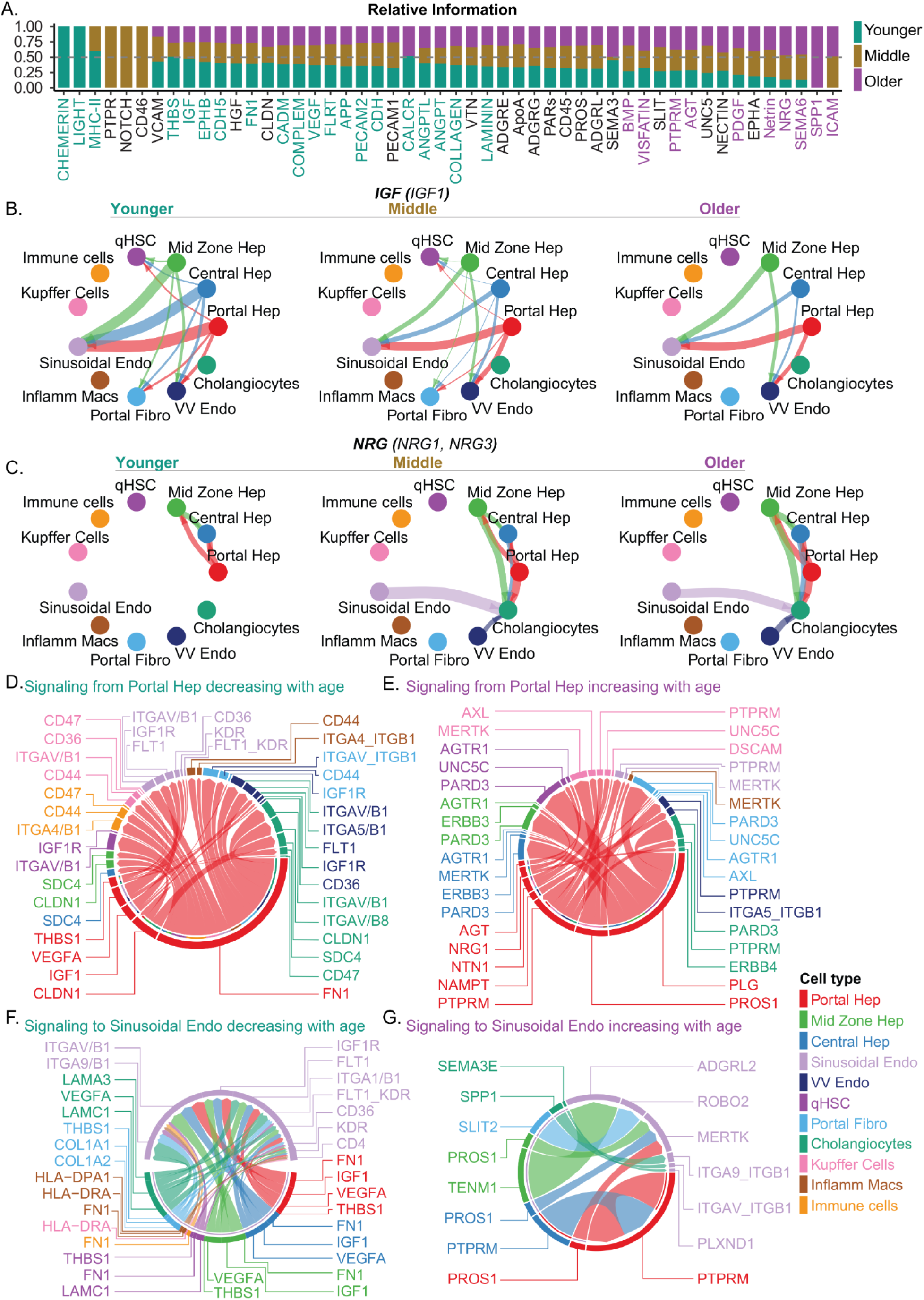
Age-related changes in signaling pathways and ligand-receptor interactions. **(A)** Signaling pathways (p ≤ 0.05) were ranked according to differences in overall information flow within inferred networks across younger, middle, and older age groups. Pathways labeled in purple are more enriched in older age groups. Pathways labeled in teal are more enriched in younger individuals. Pathways labeled in black show no significant enrichment between younger and older cells. **(B)** The chord circle plot illustrates IGF signaling pathways with enriched ligand (IGF1) across cell types in younger, middle, and older populations. Strength of communication is indicated by the thickness of the line, and arrows indicate direction. **(C)** NRG signaling with enriched ligands (NRG1 and NRG3) across age groups. **(D-G)** Chord diagrams represent aging-specific ligand-receptor (L-R) interactions. Outer rings denote different cell types, while the links illustrate interactions, with ligands expressed by the source cell type and receptors expressed by the target cell type. L-R signaling originating from portal hepatocytes enriched with decreased **(D)** and increased age **(E)** L-R signaling to sinusoidal endothelial cells enriched with decreased (**F**) and increased age (**G**) age.

We then visualized cell-cell communication across cell types and subtypes with increasing age. For example, the IGF signaling pathway, enriched in younger cells (**Fig. 6B**), shows robust signaling from hepatocytes to sinusoidal and VV endothelial cells, with signaling to qHSCs and portal fibroblasts absent in older cells. In contrast, older cells show enrichment in pathways including NRG, with stronger signals between hepatocytes and new signals developing from hepatocytes to cholangiocytes, as well as from sinusoidal and VV endothelial cells to cholangiocytes compared to younger cells (**Fig. 6C**). These findings highlight how signaling pathways vary in strength and involve different cell types as the liver ages.

To better understand the individual molecules responsible for these cell-cell interactions, we next visualized L-R pairs. We focused on portal hepatocytes and sinusoidal endothelial cells (**Fig. 6D-G),** and included data for each cell type in **Table S23**. L-R pairs, including the IGF1 (ligand) expressed by portal hepatocytes and IGF1R (receptor) expressed by sinusoidal endothelial cells, VV endothelial cells, qHSCs, and portal fibroblasts are enriched in younger donors (**Fig. 6D**). In addition, the combination of VEGFA (ligand), expressed by portal hepatocytes, and FLT1 (receptor), expressed by sinusoidal and VV endothelial cells, or KDR (receptor), expressed by sinusoidal endothelial cells, are also enriched in younger donors.

With older age, enriched L-R pairs include the combination of NRG1 (ligand), expressed in portal hepatocytes, and ERBB3 (receptor), expressed in other hepatocyte populations, or ERBB4 (receptor), expressed by cholangiocytes (**Fig. 6E**). Additionally, the combination of PLG (ligand), expressed by portal hepatocytes, and PARD3 (receptor) expressed by qHSCs, portal fibroblasts, cholangiocytes, and other hepatocytes is also enriched in older donors. These results demonstrate change in L-R pairs that occur with age, including enrichment of IGF1 signaling in younger livers and NRG1 in older livers.

We observed fewer L-R pairs signaling to portal hepatocytes enriched in younger or older donors (**Fig. S15A-B)**, as compared to L-R pairs signaling from hepatocytes. In younger donors, the interactions primarily involved the combination of LAMA3 and FLRT2 (ligands) expressed in cholangiocytes and ITGA1_ITGB1 and ADGRL2 (receptors) expressed by portal hepatocytes and the combination of collagens expressed by portal fibroblasts and multiple receptors expressed by portal hepatocytes (**Fig. S15A**). Older donors showed enrichment of L-R pairs involving BMP-5/6 (ligands) expressed by portal fibroblasts and sinusoidal endothelial cells and ACVR1_BMPR2 and BMPR1A_BMPR2 receptor complexes expressed by portal hepatocytes (**Fig. S15B**).

Analysis of L-R pairs involving sinusoidal endothelial cells also displayed age-dependent shifts, with the greatest diversity observed in signals received, predominantly from hepatocytes, portal fibroblasts, qHSCs, and cholangiocytes (**Fig. 6F**, **6G, S15C-D**). In younger donors, the combination of VEGFA (ligand) expressed by hepatocytes and FLT1 and KDR (receptors) expressed by sinusoidal endothelial cells was again observed along with the combination of IGF1 (ligand) expressed by hepatocytes and IGF1R (receptor) expressed by sinusoidal endothelial cells. L-R pairs involving THBS1 (ligand) expressed by qHSCs and portal fibroblasts and CD36 (receptor) expressed by sinusoidal endothelial cells were also enriched (**Fig. 6F**).

In older donors, the signaling landscape shifted (**Fig. 6G**). The combination of SPP1 (ligand), expressed by cholangiocytes and ITGAV/9_ITGB1 (receptors) expressed by sinusoidal endothelial cells was enriched. In addition, the combinations of PROS1 (ligand) from hepatocytes along with TENM1 (ligand) from mid-zone hepatocytes and MERTK and ADGRL2 (receptors) were enriched on sinusoidal endothelial cells. The combination of SEMA3C (ligand) expressed by cholangiocytes and the receptor PLXND1 expressed by sinusoidal endothelial cells was also enriched.

We also observed age-related changes in L-R pairs with signaling from sinusoidal endothelial cells. The combination of EFNB2 (ligand) expressed by sinusoidal endothelial cells and EPHA4 (receptor) expressed by VV endothelial cells, along with ANGPT2 (ligand) expressed by sinusoidal endothelial cells and TEK and ITGA5_ITGB1 (receptors) expressed by VV endothelial cells was enriched in younger donors (**Fig. S15C**). In older donors there was an enrichment of the combination of NRG3 (ligand) expressed by sinusoidal endothelial cells and ERBB4 (receptor) expressed by cholangiocytes and the combination of ADM (ligand) expressed by sinusoidal endothelial cells and CALCRL (receptor) expressed by VV endothelial cells and qHSCs. Similarly, there was an enrichment of the combination of NAMPT (ligand) expressed by sinusoidal endothelial cells and ITGA5_ITGB1 (receptors) expressed by VV endothelial cells. (**Fig. S15D**). Together, these findings highlight L-R pairs that change with age involving portal hepatocytes and sinusoidal endothelial cells, while also cataloging the combination of L-R pairs enriched between all cell types throughout the liver.

## DISCUSSION

The liver plays a key role in numerous physiological processes. While single-cell studies have highlighted the diversity of liver cell types and disease-related changes [1–4], the influence of sex and age on healthy liver cell types remains poorly understood. Using snRNA-seq, we analyzed over 195,000 nuclei from 37 adult donors spanning seven decades of life to explore liver cell composition and heterogeneity across sex and age.

Female hepatocytes showed broader enrichment of estrogen-related and lipid–lipoprotein pathways (e.g., Estrogen Receptor/Metabolism; LDL–HDL–TG handling), consistent with higher fatty-acid oxidation and relative protection from steatosis and NAFLD [52–54]. The Ferroptosis pathway was also enriched in central hepatocytes in females, while TGF-beta signaling was more enriched in midzone and portal hepatocytes, suggesting pathways that could influence sex-linked differences in injury responses and fibrosis/repair dynamics [55–58]. In males, enrichment of chemokine signaling, glucocorticoid receptor programs, and Hippo signaling in points to pathways that may contribute to the higher burden of NAFLD/NASH and metabolic comorbidities [59–61].

To evaluate possible contributions of menopause, data were analyzed for female donors 40 years of age and younger compared to donors older than 60 [48]. While common pathways were observed in aging for both males and females, indicating general aging, not menopause, as the primary driver of changes in these pathways, the analysis also revealed pathway shifts consistent with hepatic masculinization in older females. Younger female cells were enriched for estrogen-linked programs, Estrogen Signaling (WP712), Estrogen Metabolism (WP697), Tamoxifen Metabolism (WP691), and Sulfatase and Aromatase (WP5368)—alongside Omega-9 Fatty Acid Synthesis (WP4724), Omega-3 and Omega-6 Fatty Acid Synthesis (WP4723), and Modulation Of PI3K Akt mTOR Signaling (WP5192), consistent with pre-menopausal physiology [62–64]. In contrast, older females gained male-biased programs, including Androgen Receptor network (WP2263), SREBP Signaling (WP1982), Cholesterol Biosynthesis Pathway/Hepatocytes (WP197,WP5329), and Urea Cycle and Associated Pathway (WP4595), and these pathways were all enriched in younger compared to older males (**Table S18,19,20,21**). The convergence of older female enrichments with these male-anchored axes, also supports a shift toward a male-like hepatic state as estrogen signaling declines. This interpretation aligns with evidence that declining hepatic estrogen receptor alpha (ERα) after menopause promotes transcriptomic masculinization [65] and with animal data showing higher plasma cholesterol in low-estrogen females [66].

BMP emerged as a male-enriched signaling pathway (**Fig. 3G**), driven by BMP6 expression in sinusoidal endothelial cells signaling to qHSCs, with weaker signals to hepatocytes, cholangiocytes, and portal fibroblasts[67–69]. BMP6 is induced with MASLD and has activity against inflammation and fibrosis [70], in addition to its role in regulating hepcidin (HAMP) expression in hepatocytes [71]. BMP family members also modulate DGAT2 and mTORC1 in hepatocytes to regulate lipid/triglyceride turnover [72,73] and interact with Gremlin1 to control hepatocyte senescence during progression of NAFLD/NASH [74]. Beyond iron homeostasis, BMP6 has also been shown to suppress hepatic fibrosis in NAFLD [70], highlighting its broader role in liver metabolism and pathology. Together, these findings highlight pathways by which enrichment of BMP6 signaling in males may influence susceptibility to NAFLD and related liver diseases.

Comparative analysis of protein-protein interaction networks in female and male liver cells showed enrichment of estrogen signaling networks across portal, midzone, and central hepatocytes (**Fig. 4F**, **S10**). RUNX1, KLF6, and FOS-CXLC2 form a conserved ESR1 core across all hepatocyte zones, while the ESR1 interaction with NRG1 is identified in networks for both portal and central hepatocytes. The portal network also uniquely connects ESR1 to PROX1 through DAPK1 and ESR1 to SORL1 through FAM102A. The central network uniquely connects ESR1 to IGFBP2, highlighting potential impact on lipid metabolism and hepatic steatosis [75] in central hepatocytes. While TGFBR2 is connected to ESR1 in the central and midzone hepatocyte networks (**Fig. S10**), the receptor is also enriched in female portal hepatocytes, but the interaction with SERPINE1 and LRBP1 is contained within a different cluster (**Fig. 4E**).

Machine learning analysis of gene expression in females and males, with and without X and Y chromosome genes, highlighted significant predictive power in both donor and cell type classifications. Machine learning also revealed distinct autosomal gene expression differences, including both established and previously unrecognized sex-specific differential expression (**Table S13**). *FKBP5* encodes a steroid receptor co-chaperone that interacts with glucocorticoid, estrogen, and progesterone receptors, showing sex-dependent expression influenced by hormonal environments [76,77]. Among female-biased features, A2M is estrogen-responsive and consistently expressed at higher levels in females [78,79], while PZP is a pregnancy-associated protein regulated by estrogen and progesterone[80]. PPARGC1A is a transcriptional coactivator whose sex differences are shaped by metabolic and hormonal inputs[81]. Our analysis of GTEx data further supports the conclusion that CUX2 is enriched in liver cell types in male livers (**Fig. S13, Table S14**). These findings open avenues for further research into the intricate biological pathways that govern sex differences at the cellular level.

Age-related shifts in liver cell signaling revealed that younger livers show enriched IGF and VEGFA pathways linked to growth, vascular interactions, and cellular maintenance. IGF and its receptor IGF1R prevent apoptosis, promote cell proliferation, and induce vascular endothelial growth factor (VEGF) expression [82]. Vascular endothelial growth factor A (VEGFA) regulates angiogenesis, inflammatory responses, and energy metabolism, including hepatic lipid and glycogen metabolism [83]. In contrast, older livers exhibit increased NRG1 and PLG signaling, suggesting adaptations in repair and metabolism. Neuregulins (NRGs) regulate glucose and lipid homeostasis [84] and protect against NAFLD by modulating ERBB3 phosphorylation and PI3K-AKT signaling[85]. These findings highlight age-related shifts from growth and vascular signals in younger livers to repair-focused signaling in older livers.

### Limitations

While this study provides insights into sex- and age-related differences in liver signaling pathways, limitations exist. The cohort of 37 adult donors, mostly Caucasian (35), may limit generalizability to broader populations. Samples from a single center reflect local geography, and although histologically assessed as healthy, tissues were derived from excess surgical samples, potentially not representing completely healthy livers, and we cannot rule out the potential for more subtle pathologies. Covariate-adjusted analyses were performed to control for age and sex, but the assignment of age by decade to each sample prevented exact age matching between donors. Additionally, the study lacks direct information on menopausal status among female donors. While we attempted to approximate this by comparing women ≤40 years and >60 years, this approach cannot fully capture the heterogeneity of reproductive aging and associated hormonal changes. Despite these constraints, the study highlights the broad influence of sex and age on liver cell types and subtypes.

In conclusion, our snRNA-seq analysis reveals sex- and age-related changes in gene expression, signaling, and cell-cell communication in the healthy human liver at the cell-type and subtype level. These findings offer a valuable reference for studying the impact of sex and age on liver disease progression. Future studies incorporating comprehensive clinical covariates such as BMI, metabolic status (e.g., insulin resistance indices), and circulating hormone profiles (e.g., serum estradiol) will be required to understand the influence of these factors on cellular functions within the context of sex and age.

## Declaration of AI-assisted technologies in the writing process

During the preparation of this work, the authors used ChatGPT to provide alternative versions of a small number of individual sentences from which components were selected to help improve clarity. After using this tool/service, the authors reviewed and edited the content as needed and take full responsibility for the content of the publication.

## Author contribution

R.R, J.D., and A.C.M conceived the study and designed the experiments. Samples were collected following patient consent and prepared for analysis by E.T.E, H.K., and L.F. with additional assistance from A.S., K.T., M.Q., C.F., D.B, and C.F.C. snRNA-seq was performed by L.A.-Z., S.M., and C.L. with support from Å.S., J.D., and R.J.X. Computational analyses were performed by R.R. The manuscript was written by R.R and A.C.M. with input from all authors.

## Data and materials availability

Sequencing data used for this study are included in the Gene Expression Omnibus (GEO) under accession code: GSE210077

## Acknowledgements

The authors thank Ramnik Xavier and Åsa Segerstolpe for helpful discussions, Jasneet Aneja, Helena Lau, Georg Lauer, Stefan Gentile, Carlos Fernandez-del Castillo, and Genevieve Boland for assistance in collection of liver tissue and Jonathan Eddy, Oiza Dimowo, James Cheney, and Mantero Valentino Moreno-Cheek for assistance with tissue preservation. We thank Adam Slamin, Salome Maldonado, Dan Dubinsky, and the Broad Genomics Platform for help with generation of single-nucleus RNA sequencing data. A.C.M was supported by grants from the Chan Zuckerberg Initiative. This publication is part of the Human Cell Atlas - www.humancellatlas.org/publications.

