## Supplementary_Text_Figures for "Single-cell transcriptomics reveals the impact of sex and age in the healthy human liver"

**Table of Contents**

Supplementary Materials and Methods……………………………………………….2

Supplementary Figures. …………………………………………………………..12

References……………………………………………………………………………25

### Supplementary Materials and Methods

#### Single-nucleus RNA-seq quality control and analysis

Samples were demultiplexed using either Cell Ranger[1] mkfastq (for NovaSeq 6000) or bclconvert (for NovaSeqX). The reference for alignment was created using the GENCODE v42 annotation GTF and GRCh38.p13 genome FASTA files. Sequencing reads were aligned to the reference genome (including introns) using 10x Genomics Cell Ranger v7.0.1, resulting in count matrices for each sample. Seurat (v4.3.0)[2] objects were created for each sample, requiring each gene to be detected in at least 10 cells. Cells with fewer than 500 genes detected, fewer than 500 UMIs, or more than 10% of reads mapping to mitochondrial or ribosomal genes were excluded from the analysis.

Ambient RNA contamination was corrected using SoupX version 1.6.2[3] in default mode, and potential doublets or multiplets were identified and removed with DoubletFinder v2.0.4[4], with both steps performed on a per-sample basis.

After these filtering steps, the dataset contained high-quality cells suitable for downstream analyses. Individual Seurat objects were merged and normalized using the SCTransform method in Seurat v4.3.0, with mitochondrial reads regressed out. Principal component analysis (PCA) was conducted on the normalized data using the 'RunPCA' function. Integration and batch correction across samples were performed using Harmony v1.2.0[5] with the RunHarmony function. A shared-nearest-neighbor graph was created using the FindNeighbors function, and cells were clustered with the Louvain algorithm using the FindClusters function. A resolution of 0.5 was selected, resulting in 18 clusters (0-17), chosen from a range of tested values (0.1, 0.2, 0.3, 0.4, 0.5, 0.6, 0.7, 0.8, 0.9, and 1.0). Cluster visualization in two dimensions was performed using UMAP with the `RunUMAP` function, employing the euclidean distance measure. Cell-type proportions were analyzed with the speckle R package v0.99.7[6] using the ‘propeller’ function, with cell-type annotations, sample IDs, and metadata columns (Sex or AgeGroup) as input.

**Cell annotations**

Clusters identified were assigned cell type labels using the `sctype_score` function from the ScType R package[7], utilizing the ‘ScTypeDB_full.xlsx’ database downloaded on 26/06/2023. This package assigns labels based on marker genes from the ScType database specific to the tissue of interest.

Clusters were categorized using "Liver" as the reference tissue with ScType. This resulted in three hepatic stellate cell clusters (clusters 5, 12 and 17), one (non-macrophage) immune system cluster (cluster 8), two macrophage clusters (clusters 7 and 11), three endothelial cell clusters (clusters 4, 13 and 15), one cholangiocyte cluster (cluster 10), and two unknown cluster (cluster 14 containing 1297 nuclei and cluster 16 containing 167 nuclei) that could not be assigned a specific cell type. The remaining clusters 0, 1, 2, 3, 6, and 9 were annotated as hepatocytes using ScType. Marker genes were selected through differential expression analysis for each cell type.

**Post-cluster quality control**

In the post-cluster quality control process, we utilized *MALAT1* expression as a measure of high-quality nuclei. *MALAT1*, a long noncoding RNA retained in the nucleus, is considered a marker for high-quality cells[8,9]. To identify potential low-quality cells, we examined the expression of *MALAT1* across clusters. Cells with fewer than three reads for *MALAT1* were considered as having low expression, which could indicate lower-quality nuclei. This was observed in Cluster 9 (hepatocytes with 6307 nuclei) and Cluster 14 (unknown cell type with 1297 nuclei), where the majority of nuclei showed *MALAT1* expression below this threshold (**Table S2, Fig. S3**). Therefore, we excluded these two clusters (containing 7604 nuclei) from further downstream analysis. We further excluded cluster 16 from downstream analyses, as all 167 cells originated from a single donor sample (Sample-15) (**Table S2)**.

**Cell annotation with CellTypist to validate ScType annotation**

To further validate cell type annotations from scType, we employed the CellTypist Python package (version 1.7.1)[10]. For interoperability, the Seurat object containing clustering information was converted into the h5ad format using the zellkonverter R package (version 1.8.0) and subsequently loaded into Python with scanpy (version 1.11.4)[11].

Cell type predictions were generated with the CellTypist function "annotate()" using the pretrained model Healthy_Human_Liver.pkl, which was downloaded from the CellTypist model repository. This model includes 17 curated reference cell types of human liver: B cells, Basophils, Cholangiocytes, Circulating NK/NKT, Endothelial cells, Fibroblasts, Hepatocytes, Macrophages, Mig.cDCs, Mono+mono derived cells, Neutrophils, Plasma cells, Resident NK, T cells, cDC1s, cDC2s, pDCs. Predictions were made for each cluster using the majority voting setting (majority_voting = TRUE) to ensure robust assignments.

Comparison of ScType and CellTypist predictions showed strong agreement across the major cell types (**Table S3, Fig. S4**), including cholangiocytes (cluster 10), endothelial cells (clusters 4, 13, 15), hepatocytes (clusters 0-3, 6), and macrophages (clusters 7, 11). The only differences were that hepatic stellate cells (clusters 5, 12, 17) were classified as fibroblasts, and immune system cells (cluster 8) were grouped under T cells. These differences reflect the 17 predefined reference categories in the CellTypist training set, which does not explicitly include "hepatic stellate cells" or a broad "immune system cells" category (see section "Immune system cell sub-clustering and annotation").

**Hepatocytes zonation analysis**

Hepatocytes were classified into periportal, pericentral, and intermediate zones based on the expression of key zonal markers: *ASS1*, *ASL*, *GLUL*, and *CYP2E1[12]*. To differentiate hepatocyte clusters into these zones, a Central-to-Portal Ratio (CPR) was calculated for each cluster by subtracting the summed expression of portal markers *ASS1* and *ASL* from the summed expression of central markers *GLUL* and *CYP2E1*. This CPR value serves as a quantitative measure of zonal distribution, reflecting the relative expression levels of these markers within each cluster.

Clusters were categorized into three zones based on their CPR values. Clusters with CPR values less than or equal to the lower quantile were classified as "portal" hepatocytes, representing the periportal zone. Clusters with CPR values greater than or equal to the upper quantile were classified as "central" hepatocytes, representing the pericentral zone. Clusters with CPR values falling between the lower and upper quantiles were classified as "mid-zone" hepatocytes, representing an intermediate zone. The classification process was applied to all of the hepatocyte clusters, with portal zones corresponding to Clusters 2, and 3; central zones corresponding to Clusters 0 and 6; and mid-zone hepatocytes corresponding to Cluster 1 (**Table S5**).

To assess zonal classification, we employed UMAP (Uniform Manifold Approximation and Projection) visualization, which allowed for the spatial separation of the hepatocyte zones. To assess the enrichment of portal markers (*ASS1* and *ASL*) and central markers (*GLUL* and *CYP2E1*) within the hepatocyte clusters, we generated density plots for individual genes and their joint density using the ‘do_NebulosaPlot’ function in SCpubr R package (version 2.0.0.9000). These plots allowed us to visually assess the spatial distribution of marker expression within the UMAP embedding. The joint density plots provided an integrated view of marker co-expression, offering additional validation for the zonal classification derived from the quantitative Central-to-Portal Ratio (CPR) scores. This approach highlighted the enrichment of portal and central markers in the respective clusters and aligned with the CPR-based classification, providing additional support for the zonal annotation strategy.

**Immune system cell sub-clustering and annotation**

The immune system cell population (cluster 8 with 6,544 cells) identified in the analysis was further sub-clustered from the raw data to achieve greater resolution of immune cell subtypes. Sub-clustering was performed using the same analysis pipeline described above. Cell clusters were identified by testing resolution values from 0.1 to 1.0, in increments of 0.1, and a resolution of 0.2 was selected based on the ability to define five distinct clusters (0-4). Annotation of the clusters was performed using the ScType tool with "Immune System Cells" set as the target tissue. Cluster 0, consisting of 1,969 cells, was annotated with equal scores as Memory/Effector CD4/CD8 T cells and general T cells and was renamed as T cells 1. Cluster 1, containing 1,638 cells, was annotated as Granulocytes. Cluster 2, which contained 1,116 cells, was unannotated and marked as unknown by ScType. It was also enriched in hepatocyte genes. This cluster was excluded since it was not a clear immune population. Clusters 3 and 4, containing 1,024 and 797 cells respectively, were annotated with equal scores as CD4+ and CD8+ NKT-like cells and were renamed NKT-like cells-1 and NKT-like cells-2, respectively. We then examined the expression of immune system cell marker genes from ScType across each cluster. T cell markers including *CD2*, *CD44*, and *CD69* were expressed in both Cluster 0 and Cluster 1. *CD44* and *CD69* were the only granulocyte markers from ScType expressed in Cluster 1, but these genes are also expressed in T cells. Similarly, *LCK* and *PTPRC*,  T cell markers from ScType, were expressed in Cluster 1, while *GZMA* was expressed in Clusters 3 and 4 of immune clusters. Based on these findings, we renamed Cluster 1 as a second T cell population, designated it T cells 2.

To further validate ScType cell type annotations, we employed the CellTypist Python package (version 1.7.1)[10]. For interoperability, the Seurat object containing clustering information was converted into the h5ad format using the zellkonverter R package (version 1.8.0) and subsequently loaded into Python with scanpy (version 1.11.4).

Cell type predictions were generated with the CellTypist function “annotate()” using the pretrained model “Immune_All_High.pkl”, which was downloaded from CellTypist’s model repository. This model includes 32 cell types curated reference cell types including B cells, B-cell lineage, Cycling cells, DC, DC precursor, Double-negative thymocytes, Double-positive thymocytes, ETP, Early MK, Endothelial cells, Epithelial cells, Erythrocytes, Erythroid, Fibroblasts, Granulocytes, HSC/MPP, ILC, ILC precursor, MNP, Macrophages, Mast cells, Megakaryocyte precursor, Megakaryocytes/platelets, Mono-mac, Monocyte precursor, Monocytes, Myelocytes, Plasma cells, Promyelocytes, T cells, pDC, pDC precursor. Predictions were made for each cluster using the majority voting setting (majority_voting = TRUE) to ensure robust assignments.

Comparison of ScType and CellTypist predictions showed high concordance for T cells, with clusters 0 and 1 annotated as T cells by both methods (**Table S7, Fig. S8**). In contrast, both NKT-like cell populations (clusters 3 and 4) were classified as ILC by CellTypist. These differences reflect the 32 predefined reference categories in the CellTypist training set, which does not explicitly include NKT or natural killer (NK) subpopulation.

**Differential expression and gene ontology enrichment analysis**

We conducted differential expression analysis separately for both sex and age groups to explore how gene expression patterns vary. To enhance statistical robustness and reduce the number of comparisons, we categorized our samples into distinct age brackets: younger (18-40 years), middle (41-60 years), and older (61-90 years). To further investigate aging effects within each sex, we compared younger versus older females and younger versus older males. Using Seurat version 4.3.0, we compared gene expression profiles between sexes (males vs. females) and across different age groups. Differential gene expression analysis was performed using the MAST test implemented in the 'FindMarkers' function of Seurat version 4.3.0. Donor and sex (for age group comparison) or age group (for sex comparison) were included as latent variables to control for batch effects. Genes were called significant per cell type if detected in ≥10% of cells in either group (min.pct = 0.1), showed |log2FC| ≥ 0.25, and had Bonferroni-adjusted p-values (p_val_adj) ≤ 0.1. Differentially expressed genes were identified for each cell type between sexes (males vs. females), across age groups (younger, middle, and older), and between younger and older donors within each sex.

For age-related analysis, pairwise comparisons were performed between younger and middle-aged groups, and between middle-aged and older groups, for each cell type. Relatively few genes showed statistical significance across comparisons (younger < middle < older) or (younger > middle > older) within a cell type. To define genes that increase with age (younger < middle < older) or decrease with age (younger > middle > older), we first identified statistically significant differentially expressed genes between the younger and older-aged groups. From this list, we retained genes that were followed the increasing (younger < middle < older) or decreasing (younger > middle > older) expression in the middle aged group within the same cell type. Gene ontology enrichment analysis for biological processes was performed using EnrichR version 3.2[13] library WikiPathway_2023_Human. Enrichr combined score is calculated by the logarithmic transformation of the p-value obtained from Fisher's exact test, multiplied by the z-score representing the deviation from the expected rank. Enriched categories were considered significant if they met the criteria of FDR ≤ 0.1 and contained a minimum of three genes per term (**Table S9**, **Table S10, Table S16,** and **Table S17**).

**STRING-DB network analysis**

We analyzed protein-protein interactions (PPIs) based on sex differences using the R package STRINGdb v2.10.1[14]. The STRING database (species = 9606, version = 11.5) was accessed with the network type set to full. After performing differential expression analysis, STRING was used to generate PPI networks for the upregulated genes in males and females (excluding mitochondrial and ribosomal genes), enabling the assessment of potential biological interactions between differentially expressed genes. Networks were clustered and subsequently sub-clustered using string_db$get_clusters(). For each network, up to six clusters were retained and saved for downstream visualization. Each retained cluster was then sub-clustered using the same procedure, and the results were exported. Gene ontology enrichment analysis (WikiPathways) and subcellular localization analysis was conducted using STRINGdb and networks were visualized using the STRING online server (https://version-11-5.string-db.org/ .

**Cell-cell communication analysis**

We performed cell-cell communication analysis to investigate how intercellular signaling varies by sex and age. Using CellChat v2.1.2[15], we analyzed communication patterns across sex (males vs. females) and age groups (younger: 18-40 years, middle: 41-60 years, older: 61-90 years). CellChat was applied independently to each group and then merged for comparison using the mergeCellChat function (male vs. female for sex, and younger, middle, and older groups for age). We employed key functions such as computeCommunProb, filterCommunication, computeCommunProbPathway, aggregateNet, and netAnalysis_computeCentrality with default parameters, using the CellChatDB.human ligand-receptor interaction database. Interactions where ligands were more highly expressed in the receiver cell type than in the sender were filtered.

To rank the resulting signaling networks in sex, we used the rankNet function to perform a paired Wilcoxon test (with do.stat = TRUE) in CellChat to compare differences in signaling information flow in female vs. male. This test assessed whether the overall communication probability, defined by the sum of the weights in the network, differed significantly between male vs. female for each signaling pathway. Significant pathways were ranked based on the overall information flow, with pathways enriched in males or females highlighted accordingly.

To rank the signaling networks across the age groups, we used the rankNet function in CellChat to analyze signaling information flow among younger (18–40 years), middle-aged (41–60 years), and older (61–90 years) cells. Since CellChat does not provide statistical significance for comparisons involving more than two groups, we focused on pairwise comparisons between younger and older groups to identify pathways with significant differences. These significant pathways were then highlighted in the three-group comparison to provide a comprehensive view across all age groups. These results were used to identify sex- and age-specific differences in cell-cell communication. Circle plots generated by the netVisual_aggregate function visualized communication patterns across groups.

To identify differentially regulated ligand-receptor (L-R) pairs between sexes, we used females as the positive dataset (positive fold change compared to males) in the identifyOverExpressedGenes function. L-R pairs upregulated in both sexes in different cell types were excluded.

For age-specific signaling and L-R pair analysis, we aimed to identify L-R pairs that were differentially enriched in younger or older age groups compared to the other groups. Specifically, the younger group (18-40 years) was compared against both the middle-aged (41-60 years) and older (61-90 years) groups, treating the younger group as the positive dataset in the `identifyOverExpressedGenes` function. Similarly, the older group was compared against the younger and middle-aged groups, with the older group as the positive dataset. This allowed us to identify L-R pairs uniquely enriched in either younger or older age groups. L-R pairs upregulated in both younger and older groups in different cell types were excluded.

**Nuclei sex prediction using machine learning**

The analysis of sex classification in liver cells was performed using a Random Forest classifier implemented through the ranger package version 0.16.0 to assess whether sex could be predicted based on gene expression profiles. The dataset was split into training (80%) and testing (20%) sets using the createDataPartition() function from the caret package version 6.0-93.

Two Random Forest models were trained: one using all expressed genes in the dataset, and the other excluding X and Y chromosome genes to evaluate the performance of autosomal genes alone in predicting sex. To assess whether the accuracy of sex prediction was influenced by donor age, we repeated the analysis using only autosomal genes within age groups. Donors were stratified into three age groups: younger (18-40 years), middle-aged (41-60 years), and older (>60 years). Sex prediction accuracy was then evaluated independently within each age category.

To optimize the Random Forest model, hyperparameter tuning was performed using a grid search with the parameters `num.trees` (number of trees in the forest), `mtry` (the number of features sampled for each split), and `min.node.size` (minimum number of observations in a node). The grid search considered three values for each of these parameters: num.trees (100, 200, 300), mtry (5, 10, 20), and min.node.size (1, 5, 10). The best-performing model was selected based on the highest accuracy, and the optimal hyperparameters were used to train the final model. After selecting the best hyperparameters, the final model was trained using the entire training set and then evaluated on the test set. The ROC curve and AUC were computed and visualized using the pROC package version 1.18.0. The importance of individual features in the model was assessed using the built-in feature importance functionality in the ranger package. The features were ranked by their importance in predicting sex.

#### ***CUX2* analysis**

Bulk RNA-seq data from the GTEx v8 dataset were accessed using the gtexr package (v0.2.1). The *CUX2* gene was mapped to its versioned ensemble identifier with get_genes(). Per-sample normalized expression values (transcripts per million, TPM) were obtained across all tissues using get_gene_expression(). Matching sample metadata, including tissue site and sex, were retrieved with get_sample_datasets(). To reduce noise, only tissues with a median TPM greater than one were retained for visualization. Expression distributions were plotted using ggplot2 as violin plots, stratified by sex, with overlaid boxplots to show medians and interquartile ranges.

#### **Senescence genes increasing with age**

To examine age-related changes in senescence-associated gene expression, we analyzed the dataset using the gene set defined by[16]. Gene expression was evaluated across individual cell types and subtypes to identify genes whose expression consistently increased across the three age groups (younger < middle < older). Pairwise Wilcoxon tests were performed to assess the statistical significance of differences in expression between age groups (younger vs. middle and middle vs. older). For each comparison, p-values were calculated and visualized on violin plots. A threshold of p <

0.05 was used to define significant differences, with p-values further categorized as follows: $p < 0.001$ (***), $p < 0.01$ (**), $p < 0.05$ (*), and $p \geq 0.05$ (ns, not significant).

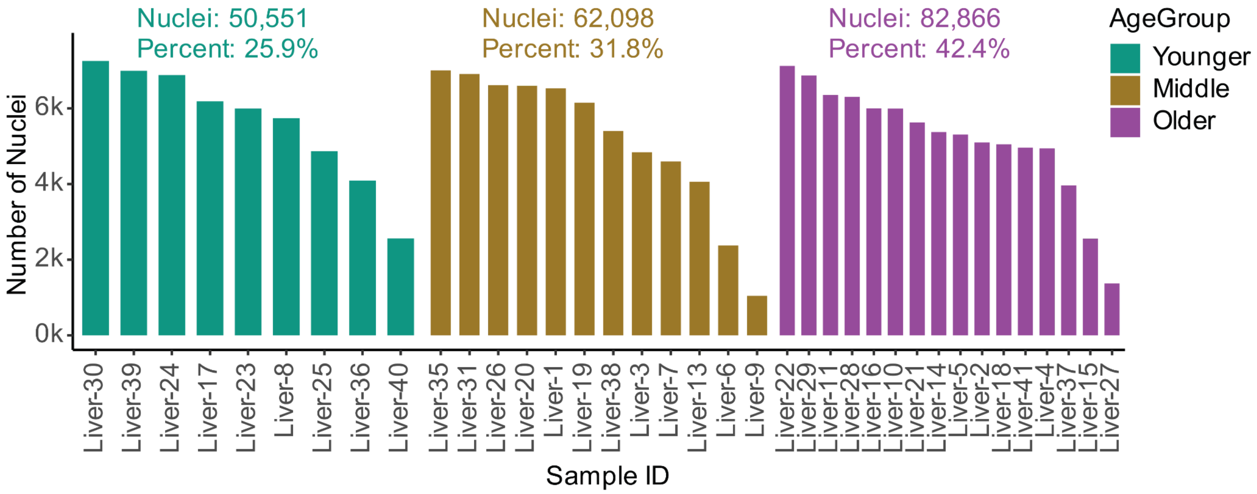

**Fig S1.** Number of nuclei in each sample, stratified by age group (younger, middle, older). Bars represent the nuclei counts within individual samples, with younger (teal), middle (mustard yellow), and older (purple) samples. This visualization provides an overview of sample composition by age group across the dataset.

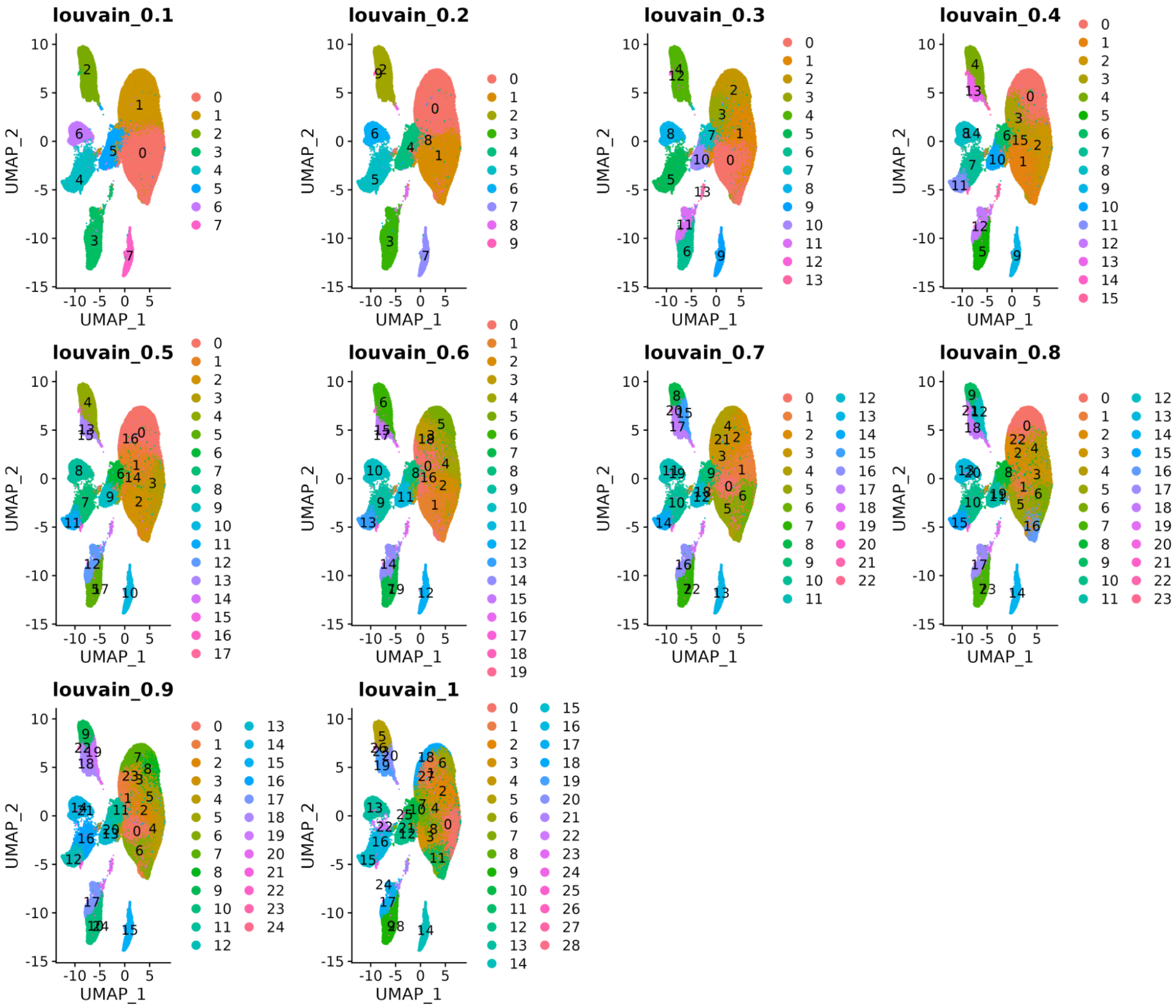

**Fig. S2**. UMAP visualization of nuclei clusters from integrated samples across resolutions ranging from 0.1 to 1.0. Each resolution highlights the clustering patterns and the corresponding number of identified clusters. A resolution of 0.5, which produced 18 clusters (0–17), was chosen for downstream analysis due to its capacity to capture biologically meaningful sub-clusters, as shown in Fig. 1C and Table S3.

A.

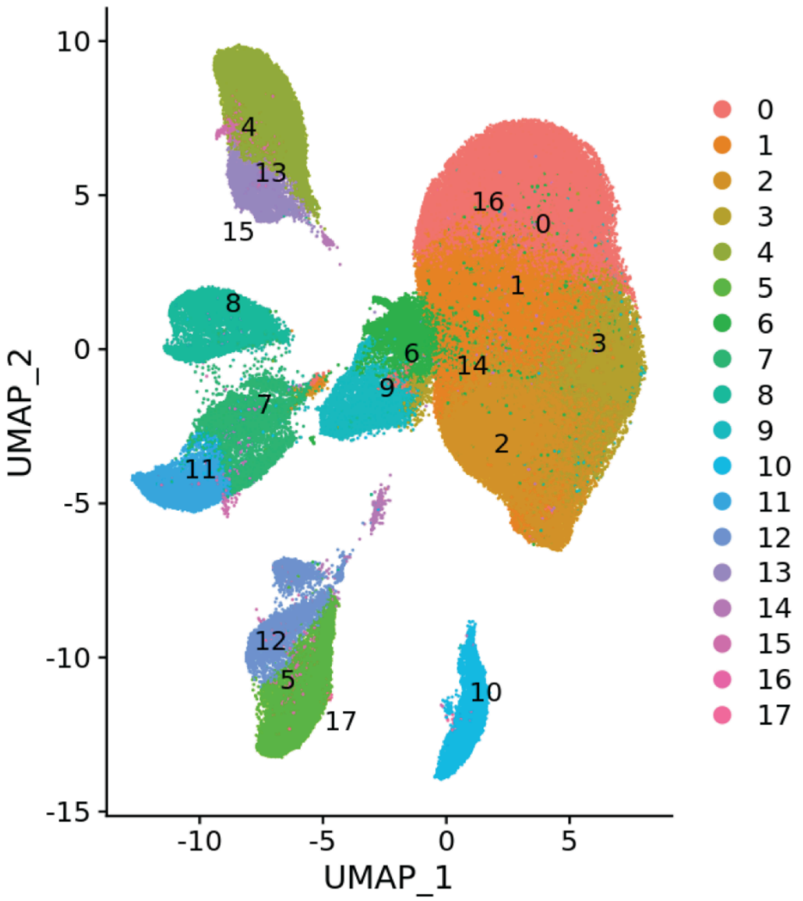
UMAP_2
UMAP_1
10
5
0
-5
-10
-15
-10
-5
0
5
4
13
15
8
7
11
6
9
14
16
0
1
3
2
12
5
17
10

B.

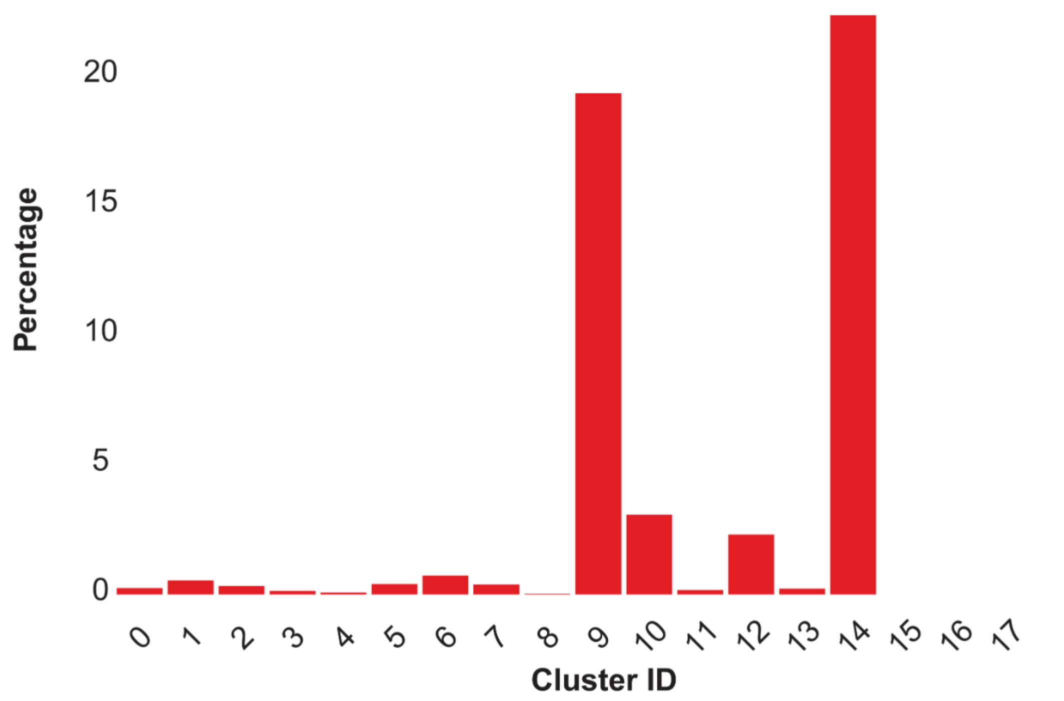
Percentage of cells with MALAT1 (Expression < 3 reads) in each Cluster
Percentage
20
15
10
5
0
Cluster ID

**Fig. S3.** (A) UMAP visualization displaying all clusters, including clusters 9 and 14, which were excluded from downstream analyses due to low *MALAT1* expression (as shown in Fig 1C). These two clusters, containing 7604 nuclei, were removed as *MALAT1* serves as a marker of nuclear quality (Clarke and Bader 2024; Montserrat-Ayuso and Esteve-Codina 2024). (B) Histogram illustrating the distribution of *MALAT1* reads per nucleus across all clusters. Clusters 9 and 14 showed a notable reduction in *MALAT1* expression, with many nuclei exhibiting fewer than three detected reads.

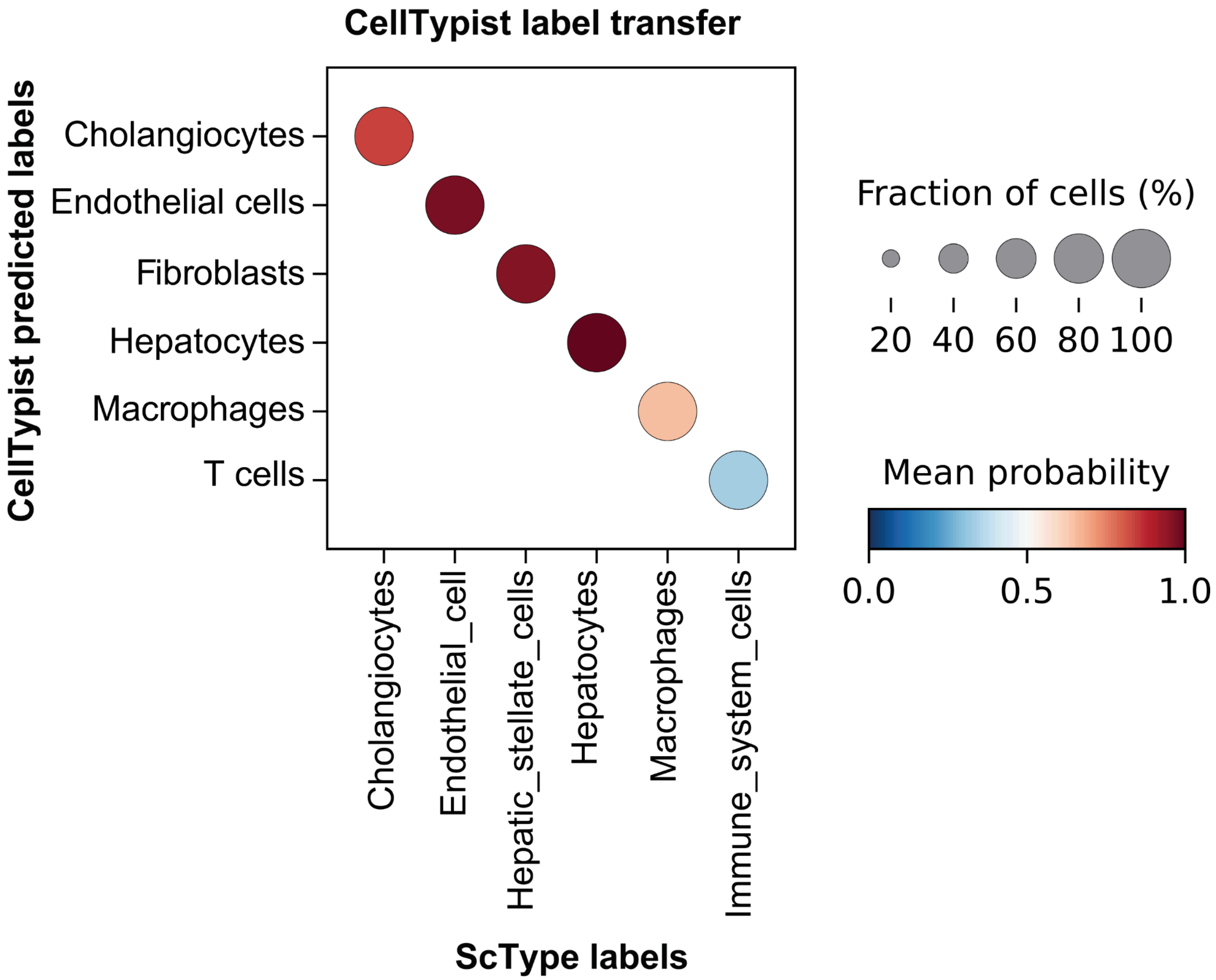

**Fig. S4. CellTypist–ScType annotation concordance**. Rows display CellTypist labels; columns display ScType labels. Bubble size represents the percentage of cells in each ScType group assigned to the corresponding CellTypist label, and bars indicate the classification probability. Comparison of ScType and CellTypist predictions shows strong agreement across major cell types (**Table S3**), including cholangiocytes (cluster 10), endothelial cells (clusters 4, 13, 15), hepatocytes (clusters 0–3, 6), and macrophages (clusters 7, 11). Hepatic stellate cells (clusters 5, 12, 17) are mapped to fibroblasts, and immune system cells (cluster 8) are mapped to T cells, reflecting the 17 predefined categories in the CellTypist training set, which do not explicitly include “hepatic stellate cells” or a broad “immune system cell” label (**Supplementary Material**).

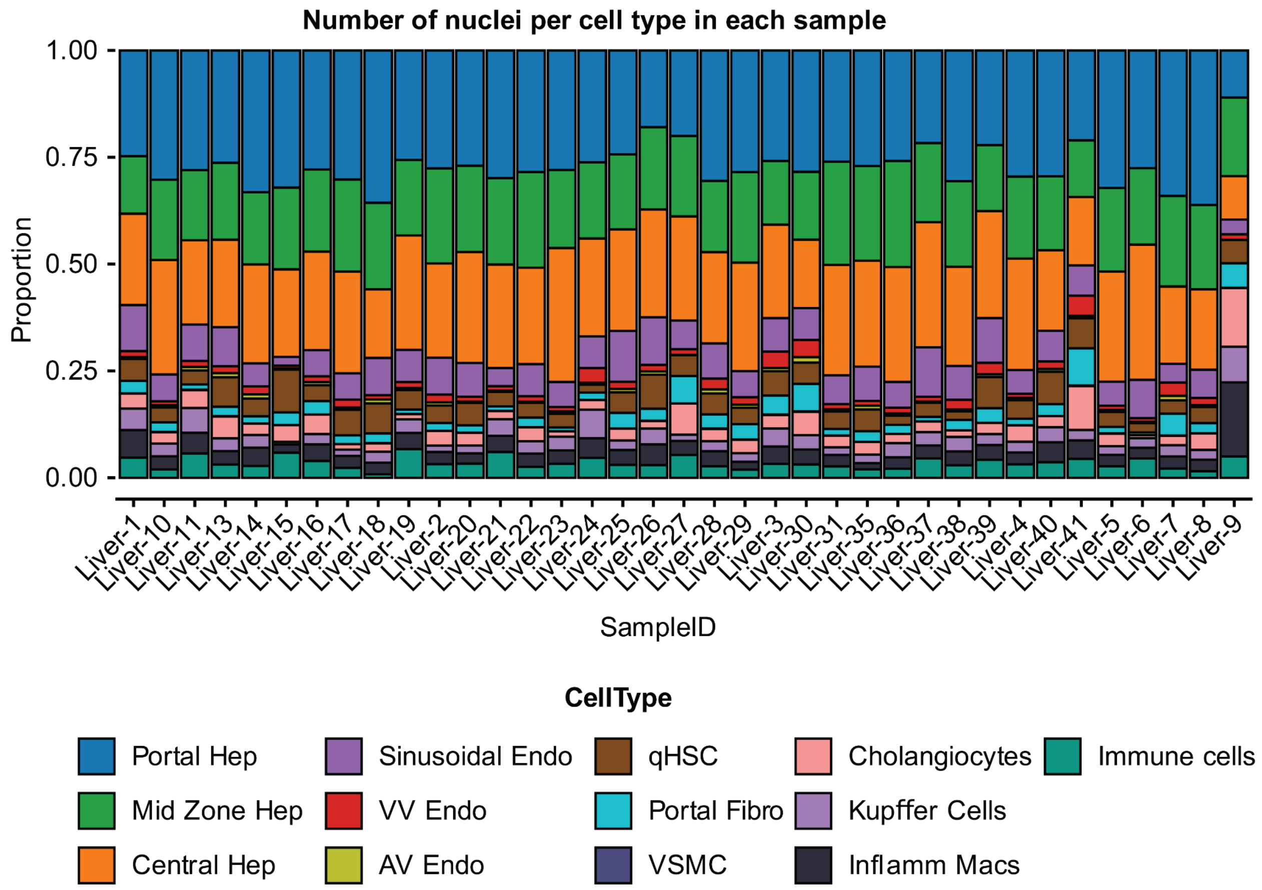

**Fig S5.** This panel illustrates the proportions of each cell type across all samples, showing a uniform representation of cell types with no sample-specific biases. The even distribution confirms that cell types are consistently represented across samples, indicating well-balanced sampling and minimal influence of technical sample artifacts.

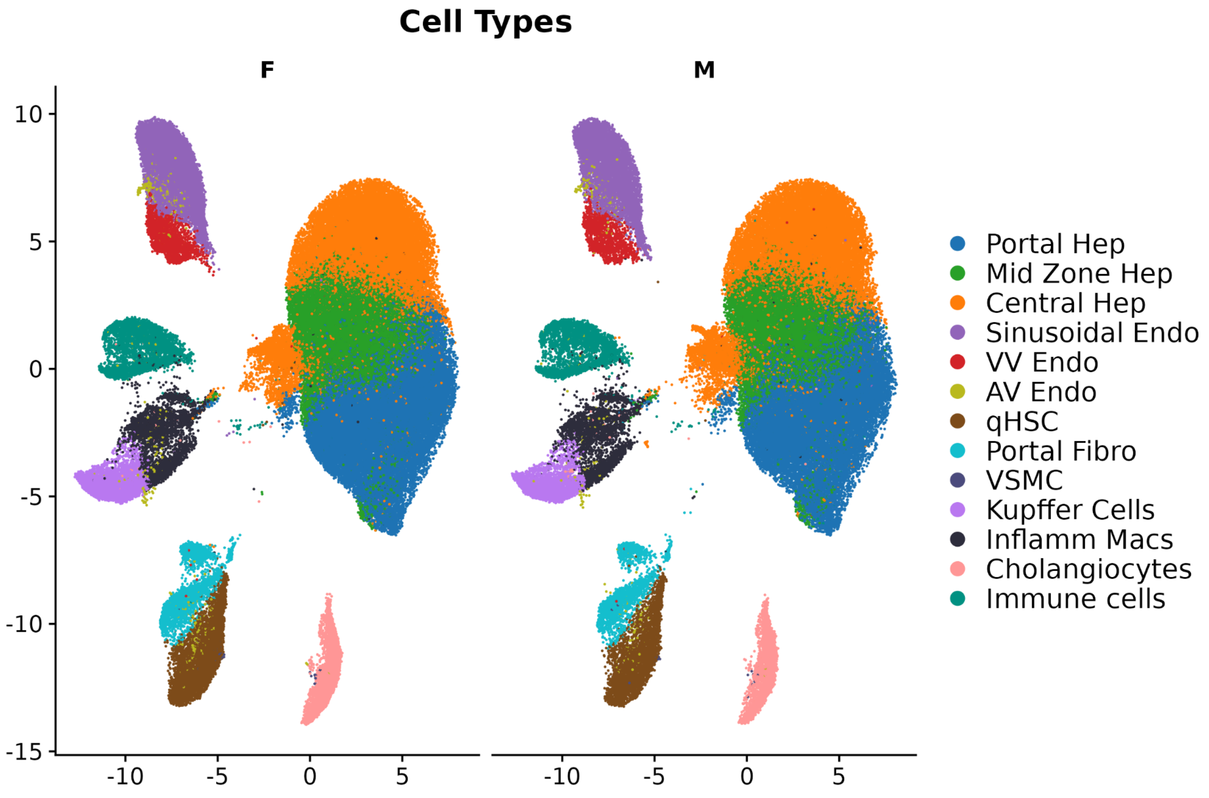

**Fig S6.** UMAP representation of cell types, split by sex, with female (F) cells on the left and male (M) cells on the right. The plot demonstrates a similar distribution of cell types across clusters, indicating comparable patterns of cell type composition between sexes.

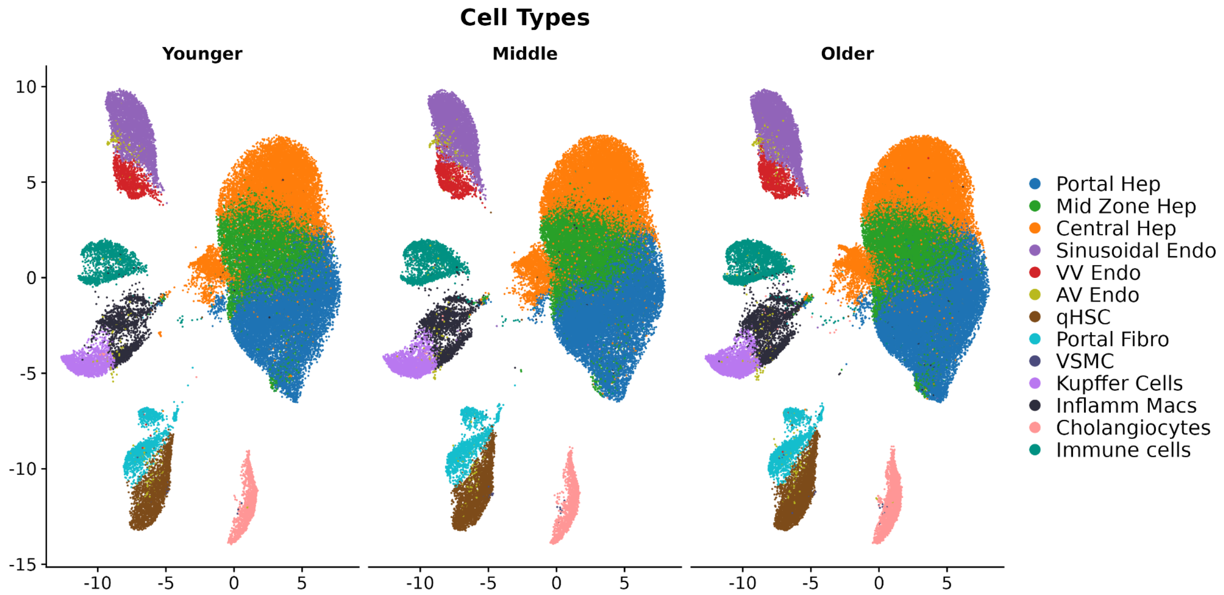

**Fig S7.** UMAP representation of cell types, split by age group, with younger cells on the left, middle-aged cells in the center, and older cells on the right. The plot demonstrates a similar distribution of cell types across clusters, indicating comparable patterns of cell type composition between age groups.

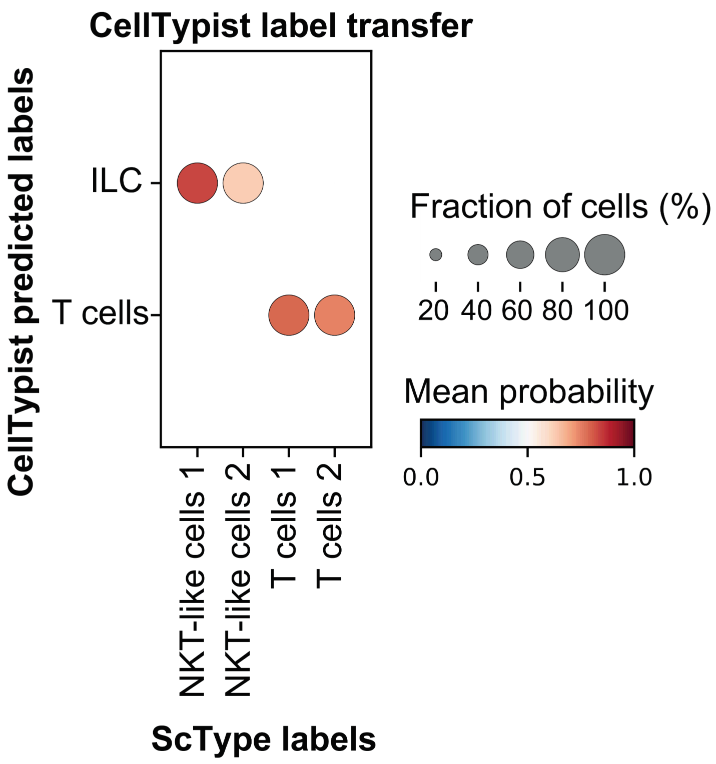

**Fig S8. CellTypist–ScType annotation concordance**. Rows display CellTypist labels; columns display ScType labels. Bubble size represents the percentage of cells in each ScType group assigned to the corresponding CellTypist label, and bars indicate the classification probability. Comparison of ScType and CellTypist predictions shows strong agreement across major cell types, including T cells, with clusters 0 and 1 annotated as T cells by both methods (**Table S7**). In contrast, both NKT-like cell populations (clusters 3 and 4) were classified as ILC by CellTypist. These differences reflect the 32 predefined reference categories in the CellTypist training set, which does not explicitly include an NK-T or natural killer subpopulation (**Supplementary Material**).

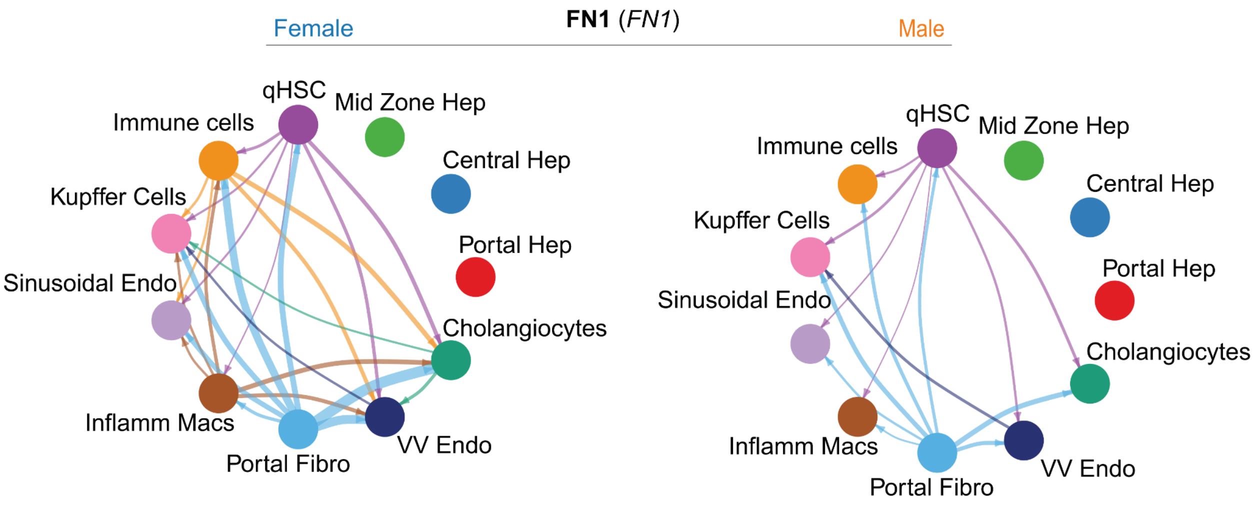

**Fig. S9.** Chord circle plots display significantly interacting pathways and communication probabilities of the FN1 signaling network, which is higher in females than males. The enriched ligand is FN1. Hepatocyte zonation groups were excluded to visualize other connections.

A. Central Hep (Female, Cluster 3)

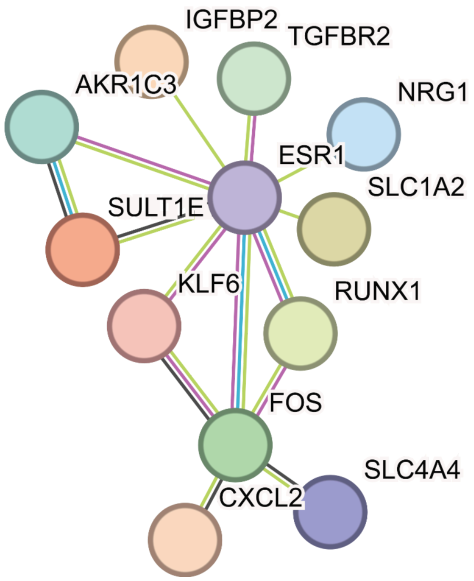

B. Mid-zone Hep (Female, Cluster 1)

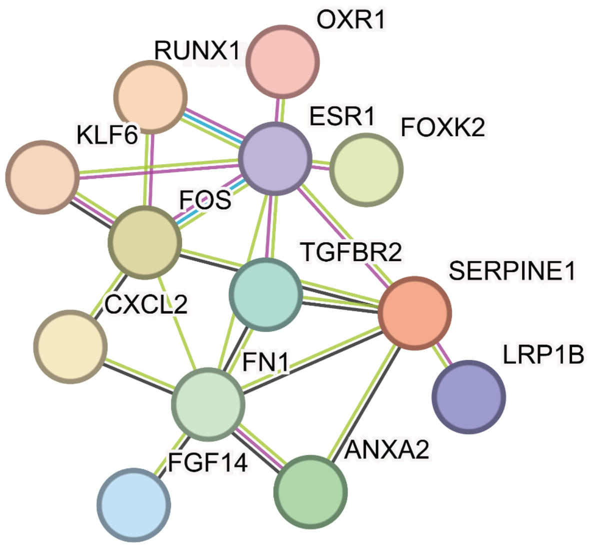

**Fig S10.** STRINGDB network analysis of protein-protein interactions (PPIs) in female liver cells. The networks visualize physical interactions between proteins based on evidence from STRINGDB. Nodes represent proteins, while edges (lines) depict interactions, with different colors indicating evidence from distinct sources. **A.** ESR1-centered gene network in central hepatocytes. **B.** ESR1-centered gene network in mid-zone hepatocytes.

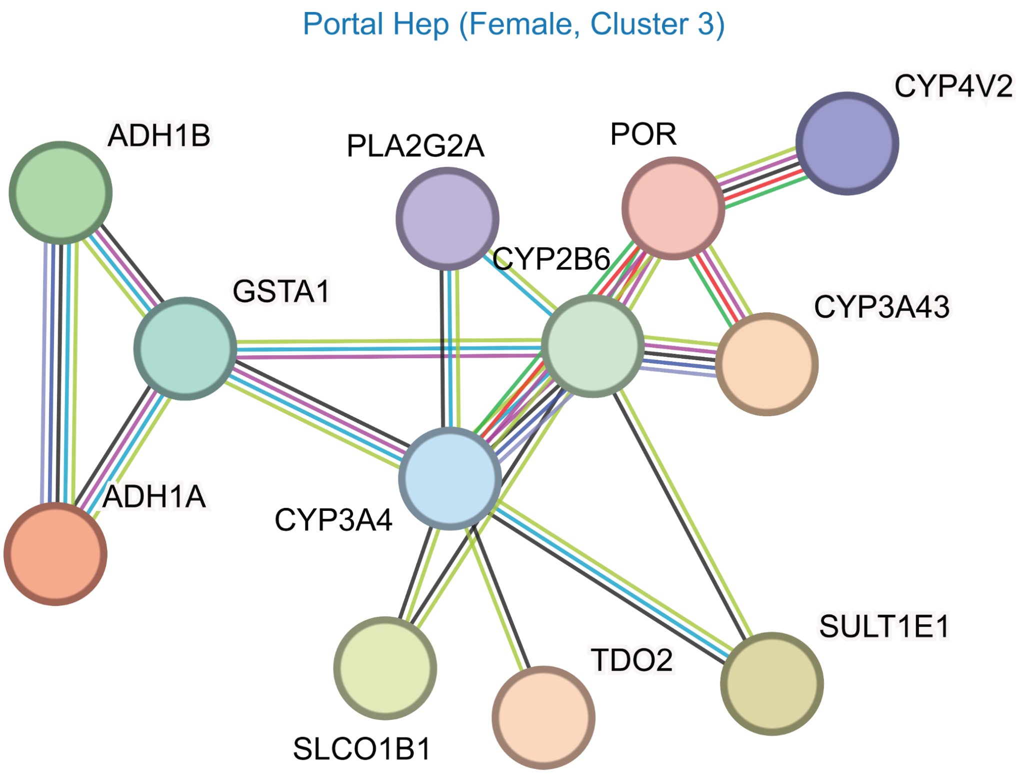

**Fig S11.** STRINGDB network analysis of protein-protein interactions (PPIs) in female liver cells. The networks visualize physical interactions between proteins based on evidence from STRINGDB. Nodes represent proteins, while edges (lines) depict interactions, with different colors indicating evidence from distinct sources.

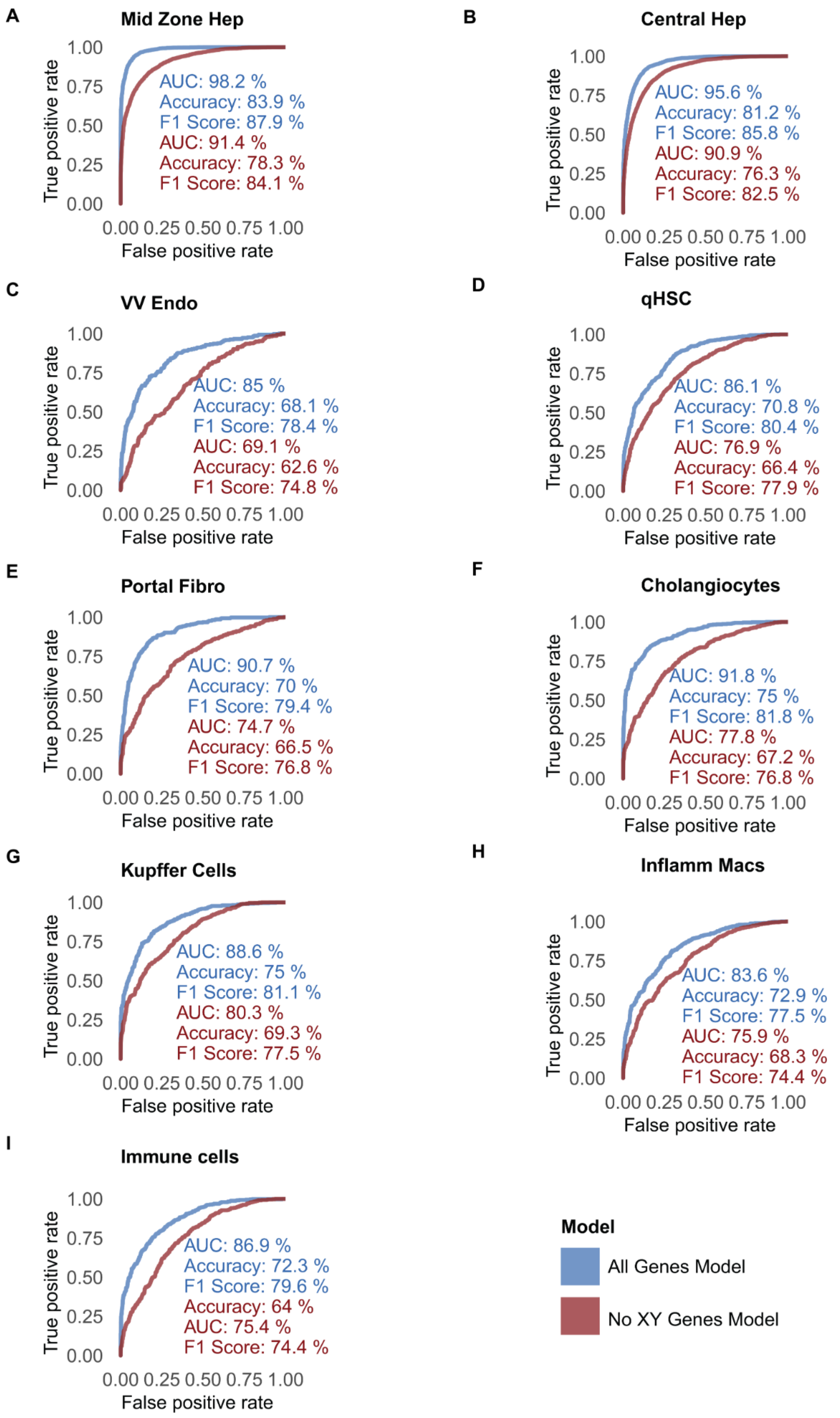
A
Mid Zone Hep
True positive rate
1.00
0.75
0.50
0.25
0.00
AUC: 98.2 %
Accuracy: 83.9 %
F1 Score: 87.9 %
AUC: 91.4 %
Accuracy: 78.3 %
F1 Score: 84.1 %
0.00 0.25 0.50 0.75 1.00
False positive rate
B
Central Hep
AUC: 95.6 %
Accuracy: 81.2 %
F1 Score: 85.8 %
AUC: 90.9 %
Accuracy: 76.3 %
F1 Score: 82.5 %
C
VV Endo
AUC: 85 %
Accuracy: 68.1 %
F1 Score: 78.4 %
AUC: 69.1 %
Accuracy: 62.6 %
F1 Score: 74.8 %
D
qHSC
AUC: 86.1 %
Accuracy: 70.8 %
F1 Score: 80.4 %
AUC: 76.9 %
Accuracy: 66.4 %
F1 Score: 77.9 %
E
Portal Fibro
AUC: 90.7 %
Accuracy: 70 %
F1 Score: 79.4 %
AUC: 74.7 %
Accuracy: 66.5 %
F1 Score: 76.8 %
F
Cholangiocytes
AUC: 91.8 %
Accuracy: 75 %
F1 Score: 81.8 %
AUC: 77.8 %
Accuracy: 67.2 %
F1 Score: 76.8 %
G
Kupffer Cells
AUC: 88.6 %
Accuracy: 75 %
F1 Score: 81.1 %
AUC: 80.3 %
Accuracy: 69.3 %
F1 Score: 77.5 %
H
Inflamm Macs
AUC: 83.6 %
Accuracy: 72.9 %
F1 Score: 77.5 %
AUC: 75.9 %
Accuracy: 68.3 %
F1 Score: 74.4 %
I
Immune cells
AUC: 86.9 %
Accuracy: 72.3 %
F1 Score: 79.6 %
Accuracy: 64 %
AUC: 75.4 %
F1 Score: 74.4 %
Model
All Genes Model
No XY Genes Model

**Fig S12A-I.** Machine learning-based prediction of nuclear sex using **(A)** mid-zone hepatocytes, **(B)** central hepatocytes, (**C)** venous vascular endothelial cells, **(D)** quiescent HSCs, **(E)** portal fibroblasts, **(F)** cholangiocytes**, (G)** Kupffer like, **(H)** inflammatory macrophages, and **(I)** Non-macrophage immune cells. The blue line indicates the AUC for models trained on all genes (including sex chromosomes), while the red line represents models trained exclusively on autosomal genes. This analysis highlights that although sex chromosome genes improve performance, autosomal genes independently capture significant sex-specific expression patterns.

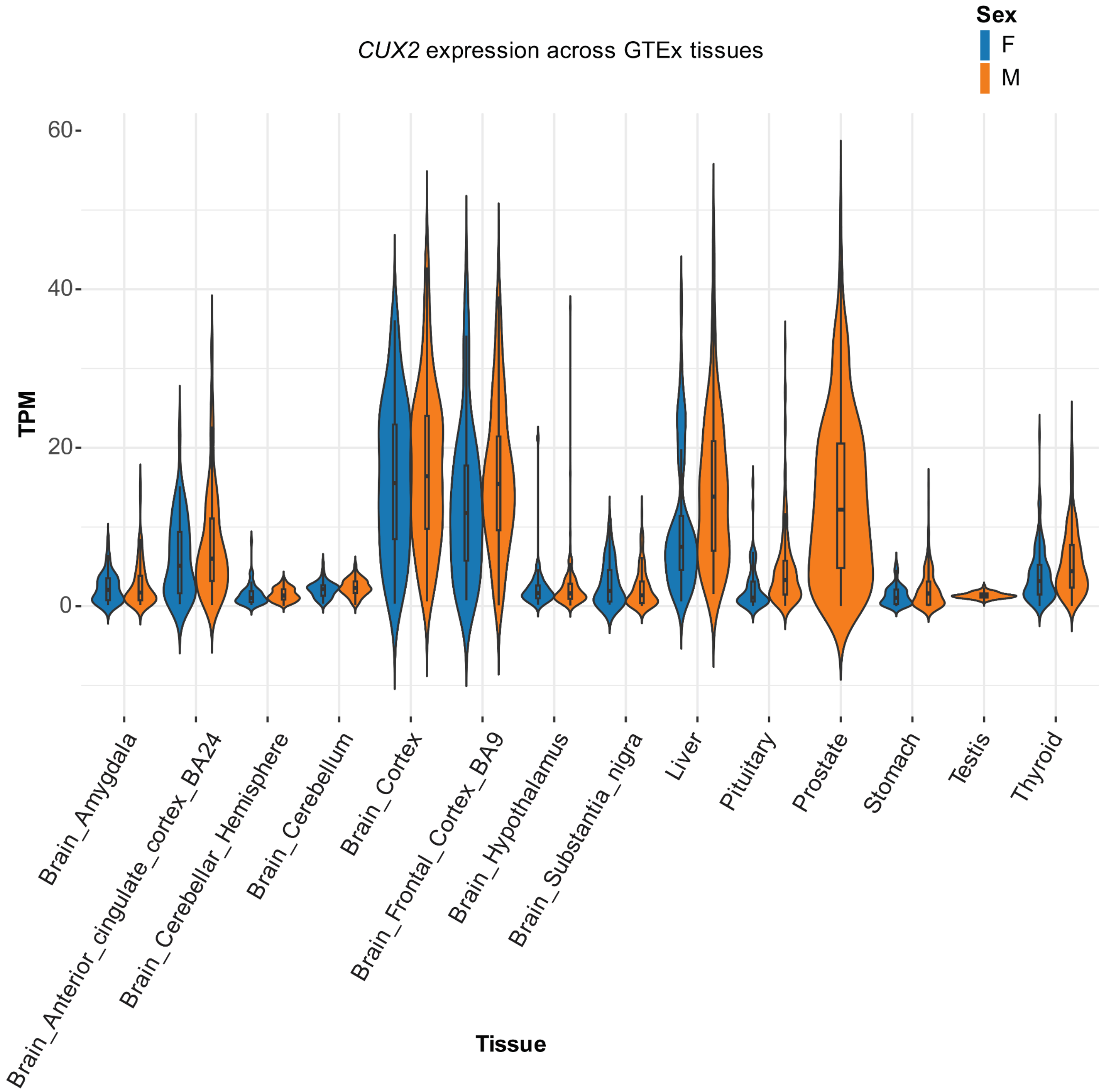

*CUX2* expression across GTEx tissues
Sex
F
M
TPM
60
40
20
0
Tissue
Brain_Amygdala
Brain_Anterior_cingulate_cortex_BA24
Brain_Cerebellar_Hemisphere
Brain_Cerebellum
Brain_Cortex
Brain_Frontal_Cortex_BA9
Brain_Hypothalamus
Brain_Substantia_nigra
Liver
Pituitary
Prostate
Stomach
Testis
Thyroid

**Fig S13. *CUX2* expression across human tissues in GTEx v8, stratified by sex.** Normalized bulk RNA-seq expression values (TPM) for *CUX2* were obtained from GTEx v8 using the gtexr R package. Tissues with a median TPM greater than one were retained for visualization. Violin plots display the distribution of expression values across all samples within each tissue, stratified by sex (blue = female, orange = male). Boxplots are overlaid to indicate the median and interquartile range. The y-axis is plotted on a linear scale (TPM), and tissue labels correspond to GTEx tissue site detail categories.

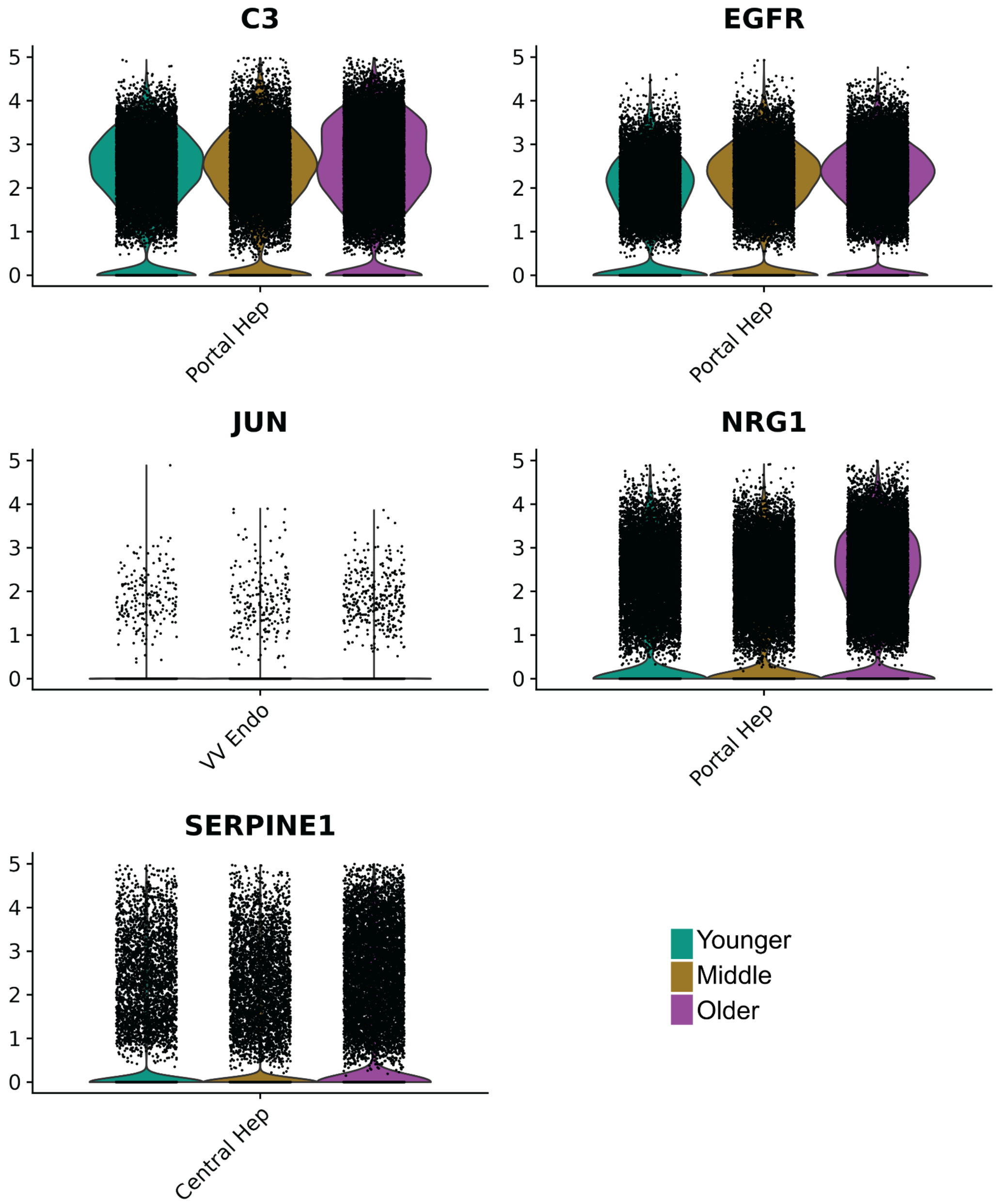

**Fig S14.** Violin plots in each panel show gene expression across three age groups: Younger, Middle, and Older. These plots highlight statistically significant changes in senescence-associated genes[16] identified in the dataset. Genes were considered significant within cell type if they were detected in ≥10% of cells in either the Younger or Older group (min.pct = 0.1), showed an absolute log2fold change (|log2FC|) ≥ 0.25, and had Bonferroni-adjusted p-values (p_val_adj) ≤ 0.1 when comparing Younger and Older groups. In addition, expression in the Middle group was required to fall between the levels observed in the Younger and Older groups.

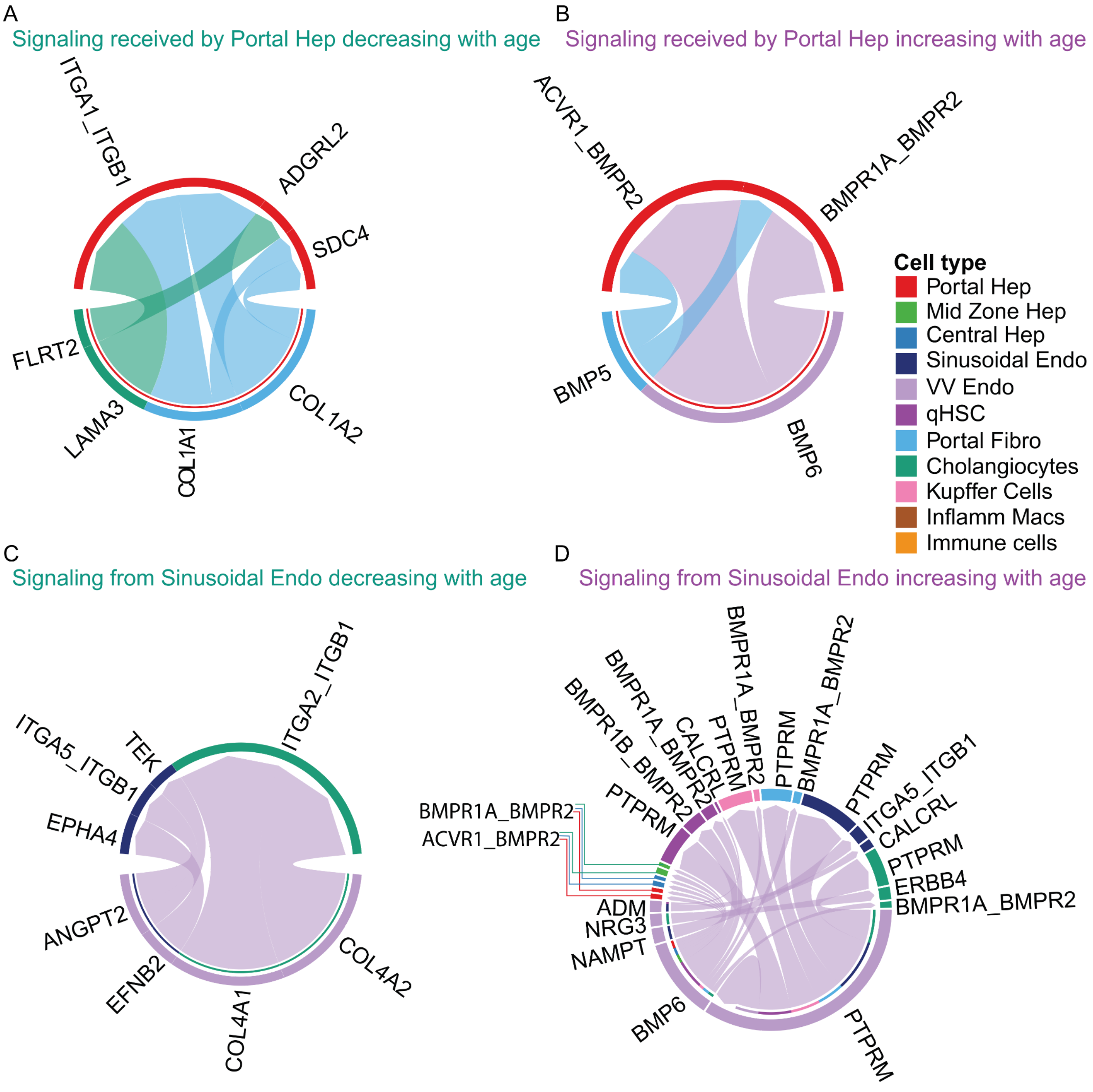

**Fig S15. A-D**. Chord diagrams representing aging-specific ligand-receptor (L-R) interactions in liver cell types and subtypes. In these diagrams, the outer rings denote different cell types, while the links illustrate interactions, with ligands expressed by the source cell type and receptors expressed by the target cell type. These visualizations highlight changes in intercellular communication networks with age. **A.** L-R signaling to portal hepatocytes decreasing with age. **B**. L-R signaling to portal hepatocytes increasing with age. **C.** L-R signaling originating from sinusoidal endothelial cells decreasing with age. **D**. L-R signaling originating from sinusoidal endothelial cells increasing with age.
